# Long-read genomes reveal pangenomic variation underlying yeast phenotypic diversity

**DOI:** 10.1101/2022.11.19.517216

**Authors:** Cory A. Weller, Ilya Andreev, Michael J. Chambers, Morgan Park, NISC Comparative Sequencing Program, Joshua S. Bloom, Meru J. Sadhu

**Affiliations:** Computational and Statistical Genomics Branch, National Human Genome Research Institute, National Institutes of Health, Bethesda, MD 20892, USA; Department of Microbiology and Immunology, Georgetown University, Washington, DC 20007, USA; NIH Intramural Sequencing Center, National Human Genome Research Institute, National Institutes of Health, Bethesda, MD 20892, USA; Department of Human Genetics, University of California, Los Angeles, Los Angeles, CA 90095, USA; Department of Biological Chemistry, University of California, Los Angeles, Los Angeles, CA 90095, USA; Howard Hughes Medical Institute, University of California, Los Angeles, Los Angeles, CA 90095, USA; Institute for Quantitative and Computational Biology, University of California, Los Angeles, Los Angeles, CA 90095, USA; Department of Computational Medicine, University of California, Los Angeles, Los Angeles, CA, 90095, USA; Department of Biology, Massachusetts Institute of Technology, Cambridge, MA 02139, USA

**Author notes:** These authors contributed equally to this work (Cory A. Weller, Ilya Andreev).

## Abstract

Understanding the genetic causes of trait variation is a primary goal of genetic research. One way that individuals can vary genetically is through the existence of variable pangenomic genes – genes that are only present in some individuals in a population. The presence or absence of entire genes could have large effects on trait variation. However, variable pangenomic genes can be missed in standard genotyping workflows, due to reliance on aligning short-read sequencing to reference genomes. A popular method for studying the genetic basis of trait variation is linkage mapping, which identifies quantitative trait loci (QTLs), regions of the genome that harbor causative genetic variants. Large-scale linkage mapping in the budding yeast *Saccharomyces cerevisiae* has found thousands of QTLs affecting myriad yeast phenotypes. To enable the resolution of QTLs caused by variable pangenomic genes, we used long-read sequencing to generate highly complete de novo assemblies of 16 diverse yeast isolates. With these assemblies we resolved growth QTLs to specific genes that are absent from the reference genome but present in the broader yeast population at appreciable frequency. Copies of genes also recombine onto chromosomes where they are absent in the reference genome, and we found that these copies generate additional QTLs whose resolution requires pangenome characterization. Our findings demonstrate the power of long-read sequencing to identify the genetic basis of trait variation.

## Introduction

The individual members of a species will have myriad trait differences, including in disease phenotypes, agricultural traits, traits that are subject to natural or sexual selection, and fundamental biological processes. Trait variation is often caused by genetic differences between individuals, and understanding how genetic variation is tied to phenotypic variation has been a central goal of genetics research since its inception under Gregor Mendel. The budding yeast *Saccharomyces cerevisiae* is a popular model organism in genetics research. It was the first eukaryotic species to have a sequenced reference genome (Goffeau et al. 1996), and, to date, most genes found in the reference genome have been functionally characterized (Engel et al. 2022). This was aided by the remarkable ease of testing the effects of genetic changes in yeast (Giaever and Nislow 2014), which has accelerated even further with CRISPR technology. Beyond the reference genome, *S. cerevisiae* presents substantial genetic and phenotypic diversity, making yeast a premier model for understanding the genetic basis of trait differences between diverse individuals. *S. cerevisiae* has been isolated from around the world and from diverse environments (Peter et al. 2018), including domestic fermentative environments, patient samples, and wild forests, especially trees in the beech family, *Fagaceae* (Lee et al. 2022). Whole-genome sequencing of diverse yeast isolates has revealed that many strains’ genomes contain genes not seen in the reference genome sequence. While the reference genome contains approximately 6000 genes, the “pangenome” comprised of all genes in the global yeast population is thought to contain approximately 5000 core genes shared across all strains and around 3000 variable genes present in subsets of strains. The presence or absence of pangenomic genes could have large effects on yeast traits. But pangenomic variation in strains’ genomes is not well annotated, as it is generally found in repetitive subtelomeric regions that recombine frequently (Louis et al. 1994) and are difficult to genotype (Yue et al. 2017). Thus, the nature of yeast pangenomic variation is still being understood.

Characterizing genome variation is essential to understanding how genetic differences cause trait variation. But there are thousands of genetic differences spread throughout genomes, including single-nucleotide polymorphisms (SNPs) and the variable presence of pangenomic genes, only some of which affect a given trait of interest. In yeast, an effective genetic strategy to locate relevant genetic differences is linkage mapping on a biparental cross (Ehrenreich et al. 2009). Two haploid yeast strains are mated to produce an F1 diploid strain, which undergoes meiosis to produce hundreds to thousands of haploid progeny strains known as segregants, whose genotypes and phenotypes are determined. If any genetic variant between the parental strains significantly affects the trait, segregants inheriting one parental genotype will have different phenotype values from segregants inheriting the other parental genotype. Thus, by partitioning segregants by parental genotype for all sites across the genome and identifying cases where the two groups’ phenotypes are significantly different, researchers can identify genomic regions, known as quantitative trait loci (QTLs), that contain genetic variants that affect the trait of interest. Importantly, linkage mapping on a biparental cross usually does not identify specific causal genetic variants, instead identifying regions in which they reside. This is because any nearby genetic variants will typically be co-inherited with the true causal variant, so partitioning segregants on the neighboring variants will register a similarly strong phenotypic difference. QTLs are thus composed of stretches of linked genetic variants and in yeast are often several kilobases (kb) in size, encompassing multiple genes, requiring further study to identify the specific causal variants they contain.

QTLs can be mapped even if the parental genome sequences are not completely known, so long as some genetic variants linked to the causal mutation are genotyped in the parents and segregants. Incomplete parental genome sequences pose a challenge, however, when closely studying QTLs to identify their causal genetic factors. To date, yeast genome sequences have predominantly been determined using high-throughput short-read DNA sequencing, which produces millions of reads, each encompassing fewer than 600 base pairs (bp) of the genome, which are then assembled into a predicted genome sequence (Peter et al. 2018). The resulting inferred genome sequences are often fragmented and incomplete, with particular assembly challenges arising from repetitive sequences and structural variants such as translocations, copy number variants, and pangenomic insertions and deletions. In contrast, the advent of long-read sequencing, which generates reads of 10 kb or longer (Koren and Phillippy 2015), allows for assembly of highly complete yeast genomes, as repetitive or diverged DNA sequences are generally short enough that a 10-kb long read can span them completely (Yue et al. 2017; Bendixsen et al. 2021; Saada et al. 2022; Donnell et al. 2022; Istace et al. 2017; Lee et al. 2022). Genome sequences assembled from long reads have the potential to significantly aid the resolution of QTLs to their true underlying causal genetic factors, especially those caused by pangenomic, repetitive, or structural variation.

A recent study performed linkage mapping on crosses between 16 diverse yeast strains (Bloom et al. 2019). Each parent strain was crossed to two other parent strains, for 16 total crosses. Approximately 900 segregants were isolated for each cross, and the segregants’ growth was measured in 38 distinct environments, including different nutritional sources, treatment with toxic compounds, and temperature and pH stress. Across the 16 biparental crosses and 38 traits, Bloom and colleagues mapped 7751 QTLs with a median size of 31.7 kb (12.7 QTLs per cross per trait), which were responsible for most of the genetic component of the phenotypic variance. Here, we present high-confidence genome sequences generated with long-read sequencing technology for the 16 parental strains. Leveraging the completeness of these genome assemblies, we identified pangenomic variation underlying major QTLs for growth on maltose, oxidative stress, sucrose, and raffinose. The causal genes we identified were unaccounted for by the reference genome, either because the genes were entirely absent, or because they were found at non-reference chromosomal locations. Our findings demonstrate the importance of pangenome characterization for resolving the genetic basis of phenotypic diversity.

## Results

### Long-read genome sequences

To determine the pangenome content of the 16 QTL-mapped strains, we generated de novo genome assemblies for each strain using Pacbio HiFi technology, which produces reads approximately 10-20 kb in length with very high sequence accuracy (Wenger et al. 2019). The final assemblies all contained precisely 17 contigs that corresponded to 16 nuclear chromosomes and the mitochondrial DNA (Supplemental Table 1). For 11 strains, every chromosome sequence started and ended with telomeric TG_1-3_ repeats, and across the 16 strains 505 of 512 non-mitochondrial contig ends had either TG_1-3_ repeats or telomere-associated Y′ or X elements (Fig. 1A) (Wellinger and Zakian 2012), indicating that the genome assemblies were largely or entirely complete.

**Figure 1:**
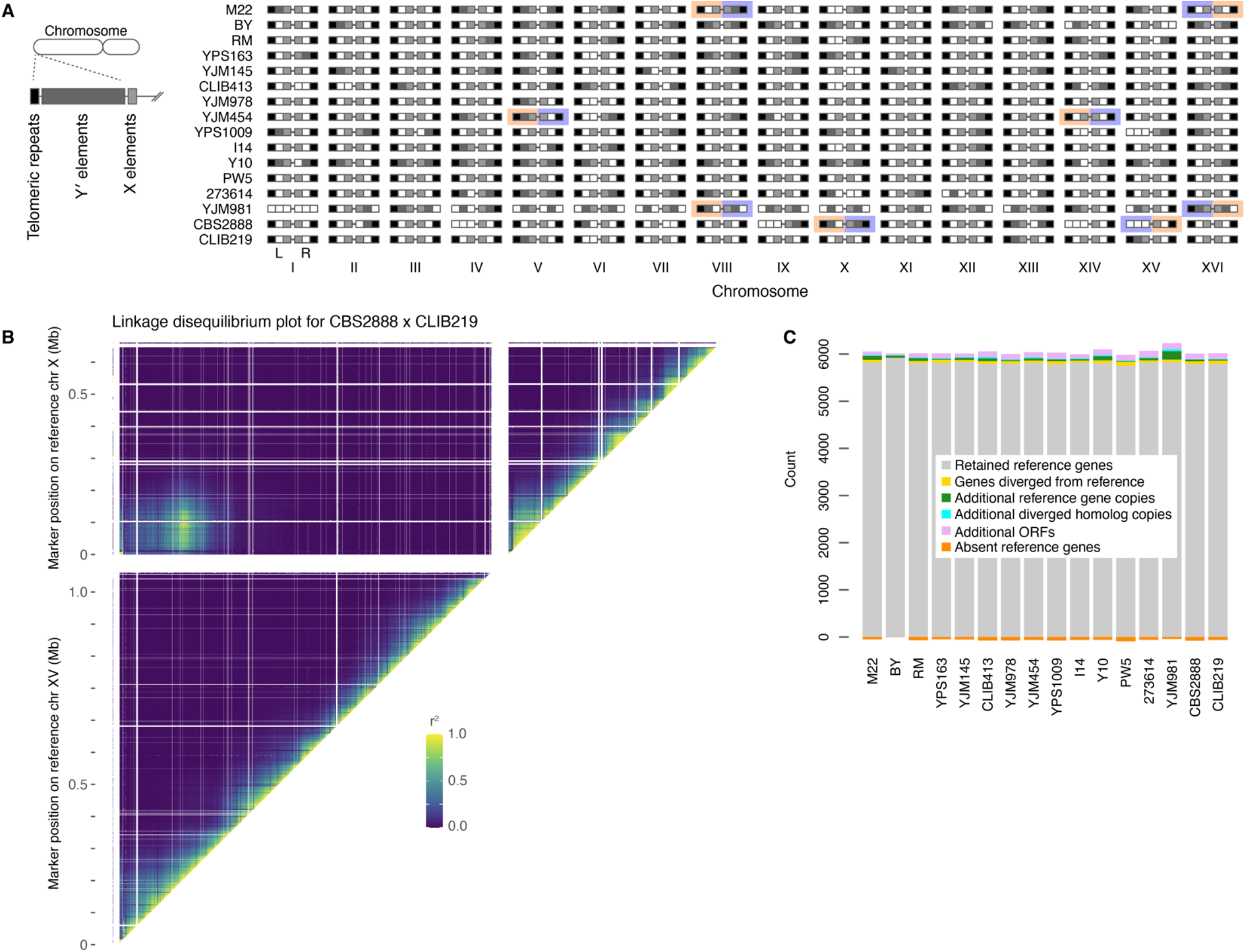
Long-read genomes. **A**. Detection of telomeric repeats and/or telomere-associated sequences at ends of assembled contigs. *S. cerevisiae* chromosomes typically end with 200-400 bp of TG_1-3_ repeats, and can also have Y′ element or X element sequences (Wellinger and Zakian 2012). The presence of TG_1-3_ repeats was called through the final 100 bp of a contig having more than 20 TG dinucleotides, while Y′ and X elements were detected by Liftoff. Dark, medium, and light gray shading indicate the presence of TG_1-3_ repeats, Y′ elements, or X elements on a given contig end, respectively, while white indicates the feature was not detected. Orange and purple highlighting in four strains indicate chromosome arms affected by reciprocal translocations. **B**. Linkage analysis of markers on reference chromosomes X and XV in the CBS2888 × CLIB219 cross. The existence of strong cross-chromosome correlation between chromosome X and XV markers is consistent with a translocation between the left arms of these chromosomes in parent CBS2888. Within chromosomes X and XV there is high correlation along the diagonal and little to no long-distance linkage, suggesting our assembled genomes accurately reflect the true chromosomal organization. **C**. Gene and ORF content in assembled genomes. Genes detected by Liftoff were classified as copies of reference genes if they had at least 95% sequence identity, and diverged homologs if they had less. Additional copies of genes were also classified as either reference copies or diverged homolog copies with a threshold of 95% sequence identity. ORFs were identified by ORFfinder and filtered to be larger than 100 codons and to not overlap other annotated genome features.

Genome assemblies of four strains had major chromosomal rearrangements relative to the reference genome (Fig. 1A). Three of these strains were previously known to have rearranged chromosomes: M22 and YJM981 have a reciprocal translocation between chromosomes VIII and XVI, and YJM454 has a reciprocal translocation between chromosomes V and XIV (Hou et al. 2014). We also detected a reciprocal translocation between chromosomes X and XV in strain CBS2888, which has not been described previously. We checked for evidence of this translocation in the genotypes of the biparental cross between CBS2888 and CLIB219 generated by Bloom et al., which resulted in 943 recombinant haploid progeny. In these segregants, we observed linkage between genetic markers corresponding to reference chromosomes X and XV, confirming the observed translocation in CBS2888 (Fig. 1B). Altogether, our assembled genomes are highly complete and reflect the true chromosomal architecture.

We annotated the genes present in the assembled genomes using a combined strategy of finding genes with homology to existing gene annotations in the S288c reference genome as well as predicting open reading frames (ORFs) de novo. The annotation tool Liftoff (Shumate and Salzberg 2021) was used to annotate genes with sequence homology to known genes in the reference genome. To capture distant homologs present in the assemblies, we set Liftoff to annotate any genes with greater than 30% nucleotide sequence identity to a known gene. To annotate any genes missed by Liftoff, such as genes with no homology to reference genes, we additionally used ORFfinder (Sayers et al. 2011) to annotate all ORFs longer than 100 codons that did not overlap any Liftoff annotations. Gene identity and content varied widely among the 16 strains. Up to 84 reference genes per strain were replaced by diverged homologs, which could reflect introgressions from other species or balancing selection within *S. cerevisiae* (Angiolo et al. 2020; Boocock et al. 2021), and up to 91 reference genes were absent without replacement (Fig. 1C). We also found as many as 141 additional ORFs as well as 216 additional copies or distant homologs of reference genes. Encouragingly, the strain BY had very few genes in these categories, as expected due to its close relation to the reference genome strain, S288c (Brachmann et al. 1998). To determine how differences in pangenome content between strains contribute to trait variation, we analyzed our annotated genomes in conjunction with previously mapped growth QTLs.

### Maltose

Maltose is a disaccharide composed of two glucose molecules. It is produced by amylase enzymes acting on starch and is a major component of brewer’s wort (He et al. 2014), which is made by enzymatic breakdown of grain and then fermented by yeast to produce beer. Metabolism of maltose in *S. cerevisiae* involves three proteins: a transporter, *MALT*, that imports maltose into the cell; a maltase enzyme, *MALS*, that converts a maltose molecule to two glucose molecules; and a transcription factor, *MALR*, that activates transcription of the *MALT* and *MALS* genes when maltose is present and glucose is absent (Brown et al. 2010). These three genes are typically found clustered together in the genome, along with *IMA* genes that encode enzymes that digest isomaltose (Teste et al. 2010). Previous analysis of diverse strains has found *MAL* loci on five different chromosomes (Charron et al. 1989), with many of these being pangenomic loci variably present between strains; for instance, the reference genome only encodes *MAL* loci on chromosomes II and VII.

Bloom et al. mapped QTLs underlying variation in the ability of the 16-isolate panel to grow on maltose. The primary genetic contributor to variable maltose growth is a recurrent QTL at the right end of chromosome VII (Fig. 2A), appearing in 12 of the 16 crosses with peak LOD scores ranging from 38 to 173. As there is a *MAL* locus in the reference genome in the subtelomeric region at the right end of chromosome VII, composed of *MAL11* (*MALT*), *MAL12* (*MALS*), and *MAL13* (*MALR*), we hypothesized that the chromosome VII maltose growth QTLs could result from variation in the functionality or presence of the *MALR, MALT*, or *MALS* genes. We examined the *MAL* genes at the chromosome VII *MAL* locus across our 16 high-quality genomes. In contrast to the reference genome, which has a single copy of each gene, the size of the chromosome VII *MAL* clusters varied dramatically from two to ten *MAL* and *IMA* genes, composed of zero to four *MALR*s, one to three *MALT*s, zero or one *MALS*s, and zero to two *IMA*s (Fig. 2B). Furthermore, the variation in *MAL* cluster size was not due to recent gene duplication events, as homologs in a cluster had low sequence identity to each other. For instance, no pair of chromosome VII *MALR*s in a single strain had higher than 85% DNA identity (Fig. 2C), with the median pair having 71% identity. Diverse pangenomic *MALR* genes have been found previously on different chromosomes in *S. cerevisiae* (Charron et al. 1989), whereas here we found that chromosome VII alone harbored homologs of all previously identified *MALR* genes except *MAL33* and YPR196W (Fig. 2C). To validate the placement of these genes on chromosome VII, we examined linkage mapping genotype data. Using previously collected Illumina short-read sequencing data for segregants in three crosses in which representatives of the different *MALR*s were segregating, we observed that segregant genotypes on the right arm of chromosome VII were strongly correlated with the presence of short reads matching the variable *MALR* sequences, confirming their assembly onto chromosome VII was correct (Supplemental Fig. 1-3). Similar to *MALR* diversity, we observe up to three distinct *MALT* genes in a single *MAL* cluster, which correspond to the previously characterized *MAL11, AGT1*, and *MTT1* transporter genes (Han et al. 1995; Dietvorst et al. 2005) and have maximum pairwise DNA identity of 91%. The sequence diversity between strains’ chromosome VII *MAL* clusters suggested the genes could have functionally diverged, which could underlie variation in growth on maltose.

**Figure 2:**
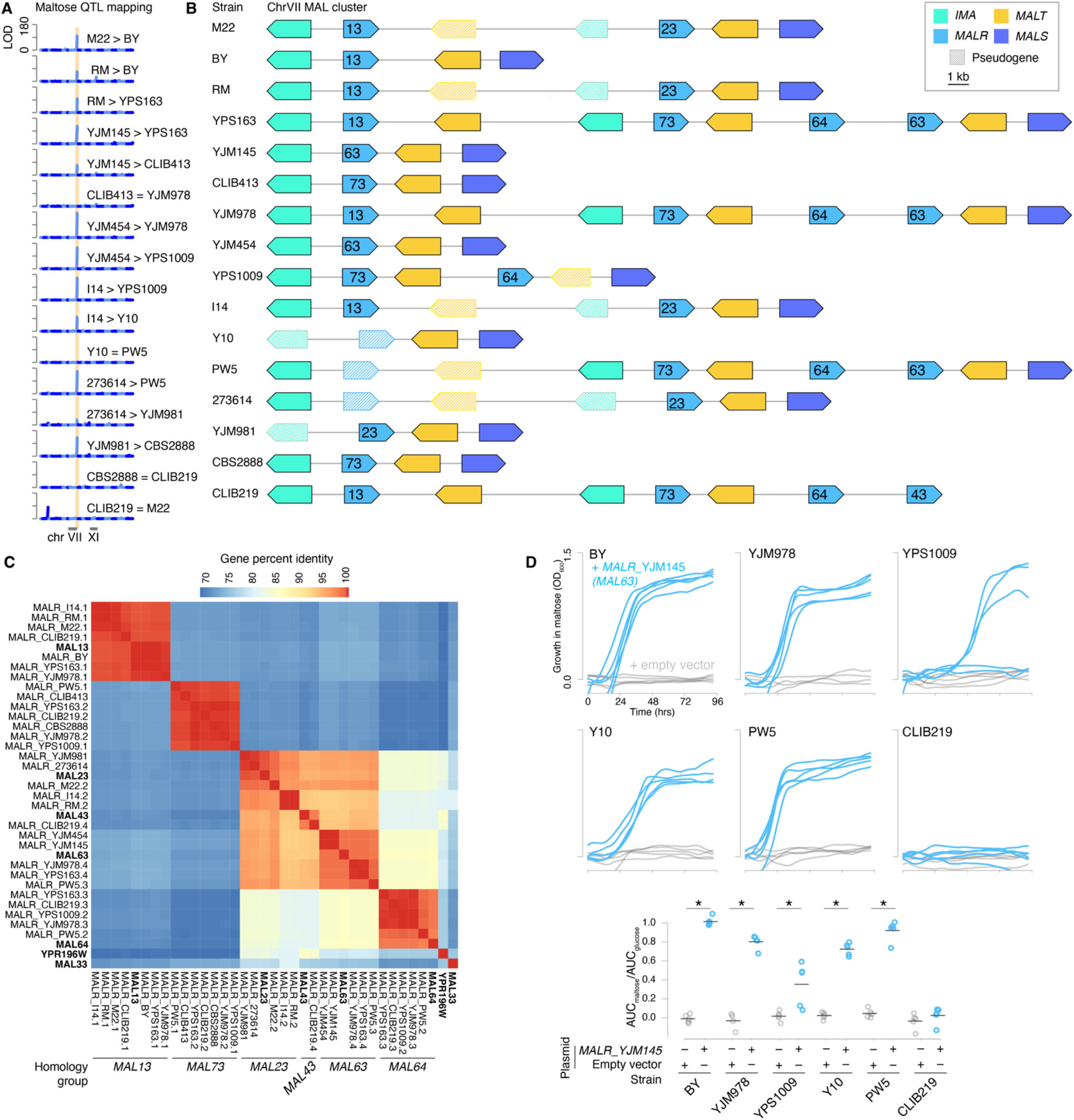
Diverse, highly complex *MAL* loci on chromosome VII. **A**. QTL mapping of maltose growth ability in 16 biparental *S. cerevisiae* crosses (Bloom et al. 2019). Each plot shows the linkage (calculated as LOD (logarithm of the odds) score), in one biparental cross, between genotype and normalized colony size after 48 hours growth on 2% maltose, plotted against genomic coordinates. Twelve of 16 crosses have a major QTL with a LOD score greater than 35 at the right end of chromosome VII, highlighted in peach; each plot is labeled with which parent provided the chromosome VII allele associated with superior maltose growth, if any. **B**. Diagram of the chromosome VII *MAL* clusters in each strain. *MALR* genes are labeled by homology group, as determined in panel C. **C**. DNA percent identity between *MALR* genes found on chromosome VII in the 16 strains, determined using Clustal Omega (McWilliam et al. 2013). Previously identified *MALR* genes are included for reference, with names in bold (Gibson et al. 1997). *MALR*s with greater than 97% sequence identity are sorted into homology groups, labeled at the bottom. As previously observed by Song et al. (Song et al. 2015), we observed a homology group of *MALR*s with less than 70.5% protein identity to any named *MALR*, which we labeled *MAL73*. **D**. (Top) Growth over time in maltose for maltose-poor strains BY, YJM978, YPS1009, Y10, PW5, and CLIB219 transformed with a plasmid expressing *MALR*_YJM145_ (blue) or empty vector (grey). *n* = 5 biological replicates, except for YJM978 + empty vector, which had 4 biological replicates. Background absorbance was subtracted, calculated from the OD_600_ (optical density at wavelength 600 nm) values of the strains grown without sugar. (Bottom) Growth was quantified as the area under the maltose growth curve normalized to the area under the glucose growth curves (Supplemental Fig. 4). Maltose growth was significantly higher for the *MALR*_YJM145_ transformants relative to empty vector for strains BY, YJM978, YPS1009, Y10, and PW5 (* p < 0.005, Tukey’s HSD following one-way ANOVA).

Previous studies examining *MALR* homologs from different *MAL* loci found that they can differ in which sugars they sense, and only some support growth on maltose. Seven distinct *S. cerevisiae MALR* genes have been previously characterized. The *MALR*s named *MAL23, MAL43*, and *MAL63* support growth on maltose, while *MAL13, MAL33, MAL64*, and YPR196W do not (Brown et al. 2010; Gibson et al. 1997). In contrast, the three *MALT* genes have all been determined to transport maltose, while varying in transport of maltotriose (Han et al. 1995; Dietvorst et al. 2005). To test whether chromosome VII *MAL* loci that do not support robust maltose growth specifically lacked a maltose-proficient *MALR*, we introduced *MALR*_YJM145_, which was associated with good maltose growth in strain YJM145 and has 99% protein identity with the known maltose-proficient Mal63, into six maltose-poor strains using a *CEN* plasmid.

We found that *MALR*_YJM145_ significantly improved maltose growth for five of the strains (Fig. 2D); the remaining strain, CLIB219, lacked *MALS* (Fig. 2B). Thus, the presence or absence of maltose-proficient *MALRs* was the major contributor to strains’ differential ability to grow on maltose.

The RM and CLIB413 genome assemblies contained *MAL* genes in the subtelomeric region on the right end of chromosome XI: *MALR* in RM and *MALR, MALT*, and *MALS* in CLIB413 (Fig. 3A, B). *MAL* genes are absent from chromosome XI in the reference genome, through the *MAL4* locus, present in some strains, is on the right arm of chromosome XI (Mortimer and Schild 1985). Linkage analysis confirmed the presence of *MALR* at the end of the right arm of chromosome XI in RM and CLIB413 (Supplemental Fig. 5, 6). The RM and CLIB413 chromosome XI *MALR* genes encoded chimeras between Mal23 and Mal43 (Fig. 3C), which are maltose proficient (Gibson et al. 1997). We thus hypothesized that these chromosome XI *MALR*s would be beneficial for maltose growth. Indeed, maltose QTLs were present at the right end of chromosome XI in all four crosses involving RM and CLIB413, with the segregants that inherited *MALR* on chromosome XI consistently exhibiting superior maltose growth (Figs. 2A, 3D, E). Combined with our observation that a maltose-proficient *MALR* was both necessary and sufficient for BY and YJM978 to grow on maltose (Fig. 2D), we concluded that the extra *MALR*s in RM and CLIB413 were responsible for the chromosome XI maltose QTLs. Notably, our high-quality genome assemblies allowed us to dissect maltose QTLs to causal underlying genes that are entirely absent from chromosome XI in the reference genome.

**Figure 3:**
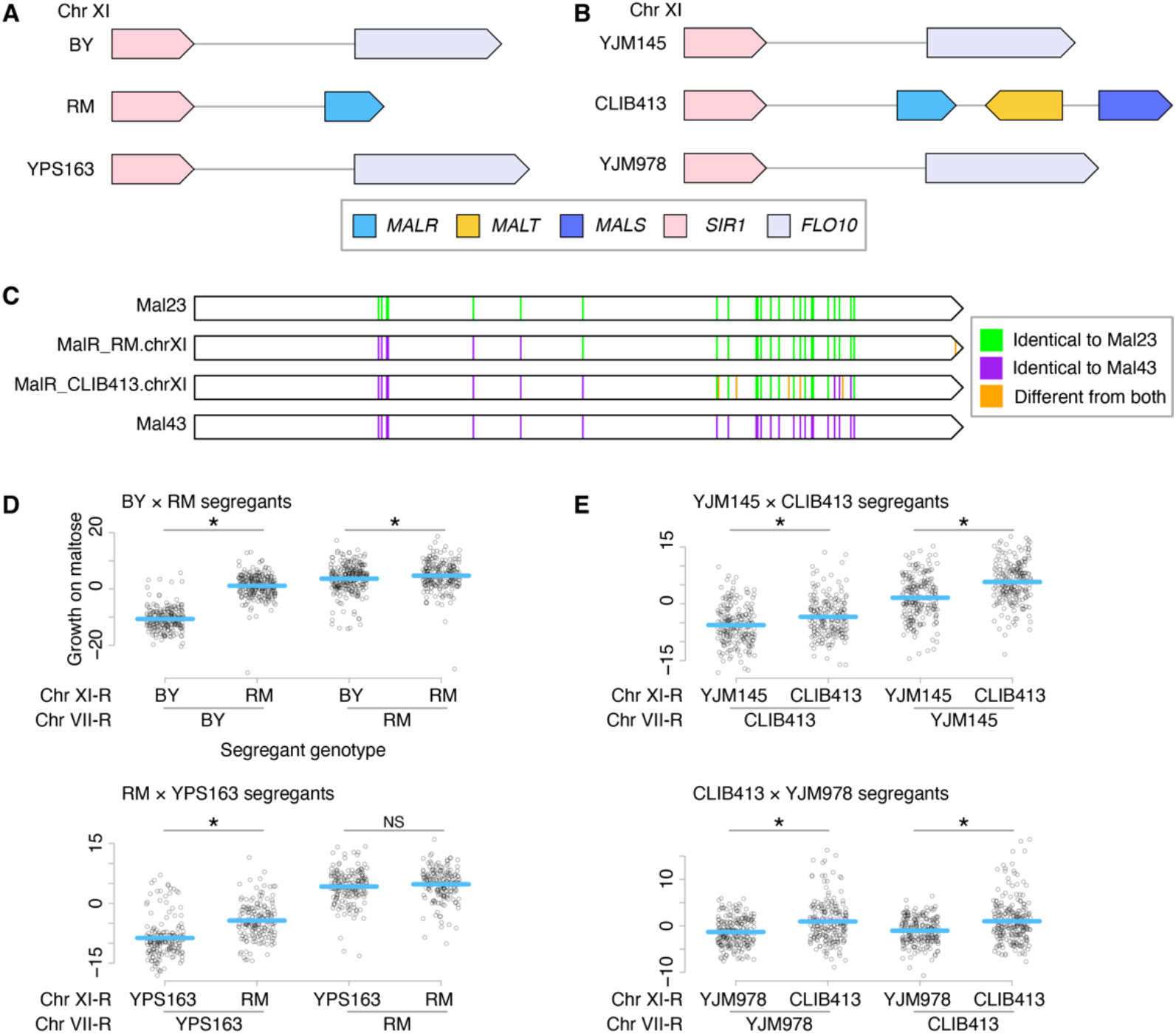
Maltose QTLs on chromosome XI are caused by non-reference *MAL* genes. **A, B**. Diagrams of *MAL* genes on chromosome XI of RM and CLIB413, along with the *MAL-*less loci in the strains they were crossed to for QTL mapping. **C**. Chromosome XI *MALR*s found in RM and CLIB413 are chimeras of *MAL43* and *MAL23*. Amino acid positions are colored green if they match Mal23 and purple if they match Mal43. Amino acid positions where either chromosome XI *MALR* differs from both Mal23 and Mal43 are colored orange. **D**. Scatterplots show the maltose growth of segregants from the BY × RM (top) and RM × YPS163 (bottom) crosses, partitioned by segregant genotype at chromosome XI-R and VII-R. Phenotypes were determined by Bloom et al. and show colony radiuses after 48 hours growth on plates containing maltose, normalized to growth on glucose plates and then scaled to have mean 0 and variance 1. The mean phenotype of each genotypic grouping of segregants is demarcated with a blue line. (NS = not significant, * p < 0.05, one-tailed Student’s *t*-test, with the alternative hypothesis that the locus with the extra *MAL* gene(s) improves growth on maltose.) **E**. Same as in part D, but for segregants from the YJM145 × CLIB413 and CLIB413 × YJM978 crosses.

### Paraquat

Paraquat is a small molecule that catalyzes the formation of reactive oxygen species, especially superoxide, from cellular electron donors (Fukushima et al. 2002). It is commonly used in the lab to study oxidative stress and is also a widely used herbicide. To explore natural variation in oxidative stress tolerance, Bloom et al. performed QTL mapping for paraquat resistance in the 16 strains.

In 11 out of 16 crosses there was a paraquat resistance QTL at the right end of chromosome XVI with LOD score greater than 10, and it was the strongest paraquat QTL for six crosses (Fig. 4A). One of the genes in the subtelomeric region at the end of chromosome XVI is *SGE1*, a transporter that extrudes cationic compounds.

**Figure 4:**
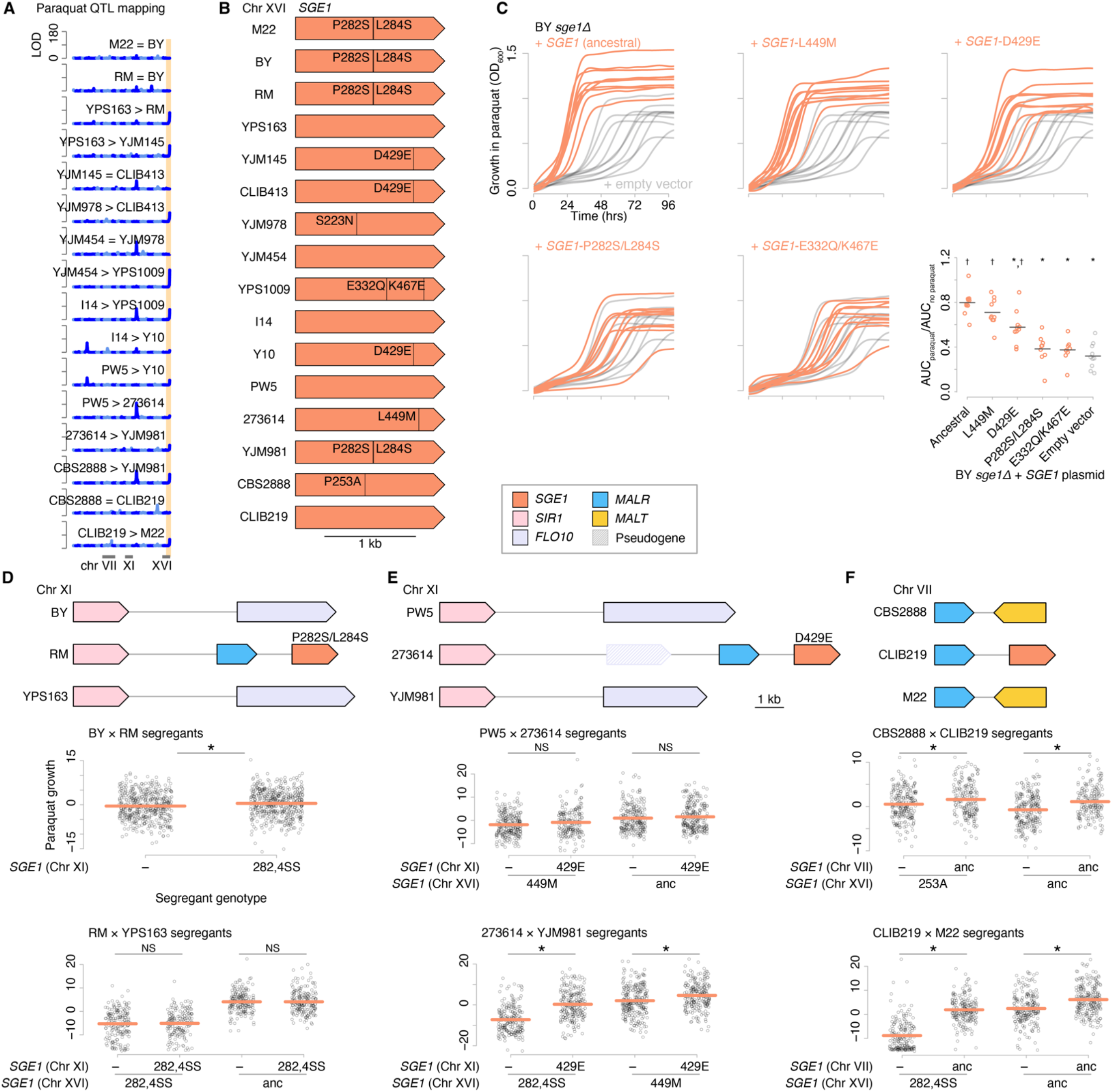
Diversity in *SGE1* underlies widespread variation in paraquat tolerance. **A**. QTL mapping of colony growth on 0.75 mM paraquat plates in 16 biparental *S. cerevisiae* crosses (Bloom et al. 2019). LOD plots are depicted as in Fig. 2A. The site of a recurrent QTL at the right end of chromosome XVI is highlighted in peach. **B**. Diagram of the *SGE1* alleles on chromosome XVI in each strain, with all derived nonsynonymous variants labeled. Variants were polarized as derived or ancestral through comparison to the *S. paradoxus SGE1* sequence. **C**. Growth over time in liquid YPD media containing 3.5 mM paraquat for BY *sge1Δ* transformed with plasmids expressing *SGE1* alleles from strains BY, YJM145, YPS1009, I14, or 273614 (orange). Growth curves are overlaid onto growth curves of BY *sge1Δ* transformed with empty vector (grey) for comparison. *n* = 10 biological replicates. Background absorbance at *t* = 0 was subtracted from all other timepoints for each curve. The last panel quantifies growth as the area under the paraquat growth curve normalized to the area under the growth curve for the same strains grown without paraquat (Supplemental Fig. 7). *SGE1* alleles from strains BY, YJM145, and YPS1009 provided significantly less protection than the ancestral allele from strain I14 (* p < 0.005, Tukey’s HSD following one-way ANOVA). In addition, the *SGE1* allele from strains YJM145, I14, and 273614 provided significantly greater protection than empty vector († p < 0.001, Tukey’s HSD), whereas the alleles from strains BY and YPS1009 did not. **D, E, F**. Additional *SGE1* genes were found in strains RM, 273614, and CLIB219 at the right end of either chromosome VII or XI, as diagrammed in D, E, and F, respectively. Scatterplots below show the growth phenotype on paraquat of each segregant in crosses involving the three strains, with segregants partitioned by *SGE1* genotype. Plots are shown as in Fig. 3, with the normalization to growth on plates without paraquat. Mean phenotypes are demarcated with orange lines. (NS = not significant, * p < 0.05, one-tailed Student’s *t*-test, with the alternative hypothesis that the extra *SGE1* copy improves growth on paraquat.)

Overexpression of *SGE1* protects against the cationic dye crystal violet (Ehrenhofer-Murray et al. 1994), and natural variation in *SGE1* between strains underlies variation in tolerance of ionic liquids used in biofuel production (Higgins et al. 2018). Paraquat is also a cation, and screens of genome-wide mutant collections have previously annotated *SGE1* overexpression as conferring paraquat resistance (Österberg et al. 2006) and *SGE1* deletion as conferring paraquat sensitivity (Helsen et al. 2020). In our genome sequences we observed that paraquat-sensitive chromosome XVI loci had one of three *SGE1* alleles: *SGE1*-P282S/L284S in strains M22, BY, RM, and YJM981; *SGE1*-D429E in strains YJM145, CLIB413, and Y10; or *SGE1*-E332Q/K467E in strain YPS1009 (Fig. 4B). We tested the effect of these three *SGE1* alleles on paraquat resistance in a BY *sge1Δ* strain, along with the ancestral allele and *SGE1*-L449M from strain 273614 (Fig. 4C). The results mirrored the QTL mapping results: the three *SGE1* alleles from paraquat-sensitive strains provided significantly less paraquat resistance than the ancestral *SGE1. SGE1*-P282S/L284S and *SGE1*-E332Q/K467E were not distinguishable from empty vector, while *SGE1-*D429E had an intermediate phenotype between empty vector and the ancestral *SGE1* (Fig. 4C). Thus, we concluded that widespread genetic variation in *SGE1* on chromosome XVI is a major contributor to differences in paraquat resistance in the *S. cerevisiae* population.

Though *SGE1* is found in the reference genome only on chromosome XVI, our genome sequencing uncovered additional copies of *SGE1* at other chromosome ends in three strains: *SGE1*-P282S/L284S in strain RM on chromosome XI, *SGE1*-D429E in strain 273614 on chromosome XI, and an additional ancestral *SGE1* in strain CLIB219 on chromosome VII (Fig. 4D-F). Linkage analysis indicated these *SGE1* copies were correctly placed on chromosomes VII and XI (Supplemental Fig. 8-10). In several cases, these extra *SGE1* loci coincided with QTLs for paraquat resistance. We examined the paraquat growth phenotypes of segregants of crosses in which these extra *SGE1* copies were segregating. Segregants inheriting the RM chromosome XI locus containing *SGE1*-P282S/L284S were slightly more paraquat-resistant than those inheriting the *SGE1-*less chromosome XI from BY, while there was no detectable benefit relative to the *SGE1-*less chromosome XI from YPS163 (Fig. 4D). There was moderate benefit of the chromosome XI locus containing *SGE1-*D429E from strain 273614, especially for segregants whose chromosome XVI locus had the inferior *SGE1*-P282S/L284S allele from YJM981 (Fig. 4E). Finally, there was a large benefit for all segregants that inherited the chromosome VII locus from CLIB219 containing the ancestral *SGE1* (Fig. 4F). These results mirrored our *SGE1* allele replacement results, in which BY *sge1Δ* grew no better with *SGE1*-P282S/L284S than with empty vector, somewhat better with *SGE1*-D429E, and best with the ancestral *SGE1*. Thus, we concluded that, through duplication, *SGE1* is additionally responsible for QTLs found on chromosomes other than where it is normally known to reside. As with the maltose chromosome XI QTLs, resolving these QTLs would not have been possible without characterization of pangenomic variation.

### Sucrose

Sucrose, a disaccharide of glucose and fructose, is a common plant sugar found in fruits and tree sap (Liu et al. 2006; Fink et al. 2018). Metabolism of sucrose in yeast requires a single protein, the enzyme invertase (Marques et al. 2016). Invertase is expressed under low glucose conditions, after which it is secreted and trapped in the yeast cell wall (Sainz-Polo et al. 2013). There, it digests sucrose to glucose and fructose, which are imported into the cell to enter standard carbon metabolism pathways.

Invertase is encoded in the *S. cerevisiae* reference genome by the *SUC2* locus on chromosome IX. Some strains carry additional genes encoding invertase close to their telomeres, with nine subtelomeric *SUC* loci having been previously identified (Carlson et al. 1985; Naumov and Naumova 2010a, 2010b), all absent from the reference genome. Across our 16 genomes, we found *SUC* genes on eight chromosomes (Fig. 5A), including three chromosome ends on which *SUC* genes have not previously been identified: chromosome I-R and chromosome III-L in strain Y10, and chromosome IX-L in strain YJM454. Linkage data supported the placement of the novel *SUC* loci (Supplemental Fig. 11-13). Despite being found on different chromosomes, most of the subtelomeric *SUC* genes have very similar sequences (Naumova et al. 2014), with 11 subtelomeric *SUC* genes found across seven chromosomes falling into a homology group with greater than 99.5% nucleotide sequence identity. This indicates they derive from a common progenitor *SUC* gene that recently spread across chromosomes (Fig. 5B), likely through inter-telomeric recombination (Carlson et al. 1985). In contrast, these subtelomeric *SUC* genes are much more diverged from *SUC2*. Their sequence identity with the reference *SUC2* allele is 93%, but this is inflated by a few regions of near-perfect identity that likely reflect recent recombination events with *SUC2* (Fig. 5C). The sequence divergence is highest in two segments, which we infer correspond to the oldest (that is, least recently recombined) regions of the subtelomeric *SUC* progenitor: nucleotides 1-432 in the catalytic domain have 88% sequence identity to *SUC2* and nucleotides 1053-1292 in the beta-sandwich domain have 81.7% sequence identity. This is higher divergence than between *SUC2* from *S. cerevisiae* and its sister species *S. paradoxus*, which have sequence identity of 92.6% and 86.3% in these regions (Fig. 5C), suggesting the original split between *SUC2* and the subtelomeric *SUC* progenitor occurred long ago, possibly prior to the speciation event from *S. paradoxus*.

**Figure 5:**
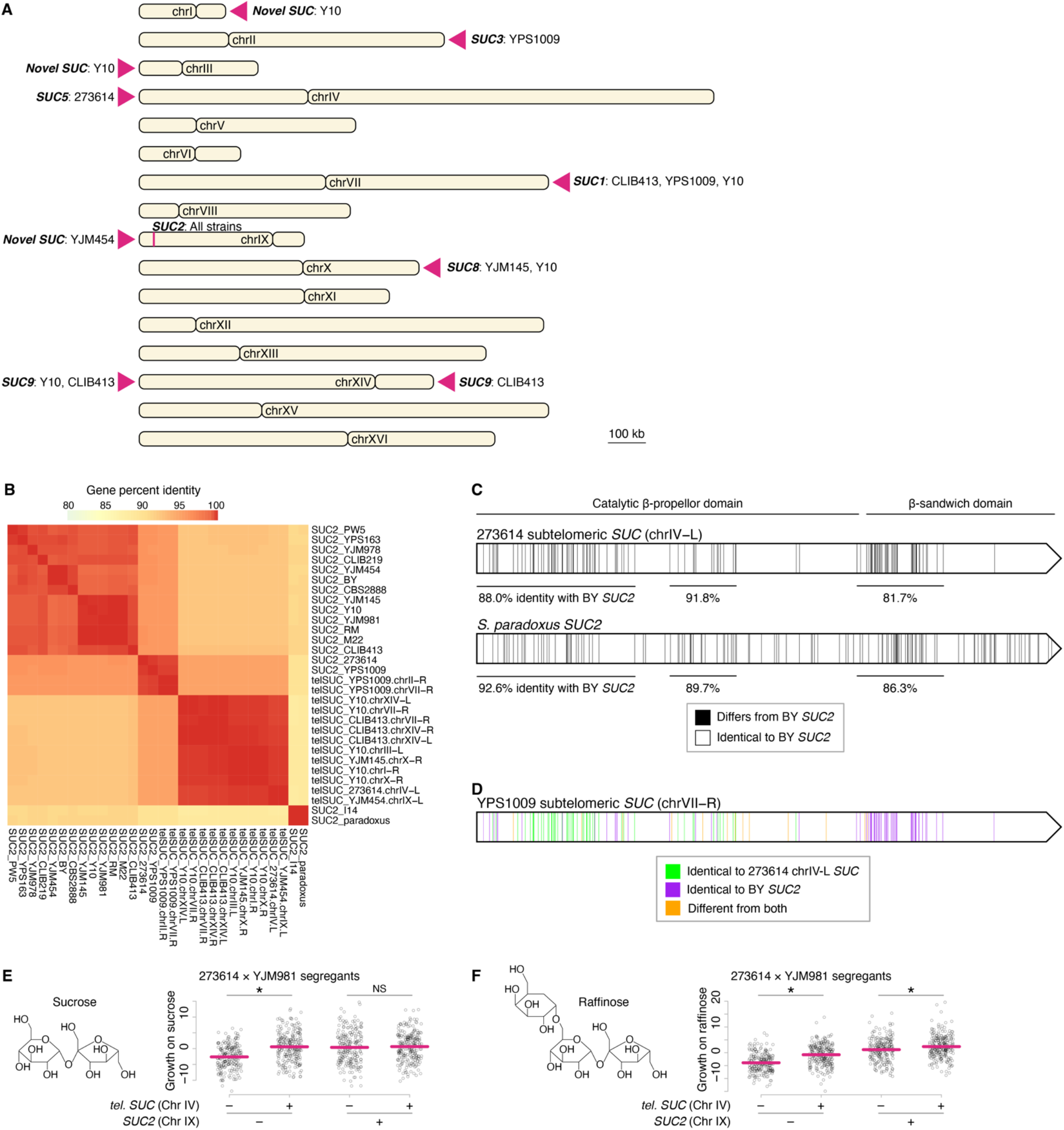
Subtelomeric copies of genes encoding invertase. **A**. Loci harboring invertase genes across the 16 genomes. Subtelomeric *SUC* loci are demarcated with magenta triangles, while the *SUC2* locus on chromosome IX is demarcated with a vertical magenta line. The strains among our 16-isolate panel carrying an invertase gene at each locus are listed along with the name of any *SUC* gene previously identified on that chromosome. The *SUC9* locus was identified on chromosome XIV (Naumov and Naumova 2010b), but whether *SUC9* resides on the left or right arm was not determined, so we labeled our identified *SUC* genes on both arms as *SUC9*. We did not identify any strains with *SUC* genes on chromosome VIII, XIII, or XVI, which have previously been found to harbor *SUC7, SUC4*, and *SUC10*, respectively (Naumova et al. 2014). **B**. Gene percent identity between *SUC* genes found in the 16 strains, along with the *S. paradoxus SUC2* reference allele, included due to its homology to *SUC2* from strain I14. **C**. Diagram of the alignment between the representative subtelomeric *SUC* gene from strain 273614 and *SUC2* from strain BY, with the positions of non-aligning nucleotides shown as vertical black lines. Three diverged regions that account for most of the non-aligning nucleotides are labeled with their percent identities to *SUC2*. **D**. A few strains carry a *SUC* allele that was formed through recombination of the predominant *SUC2* and subtelomeric *SUC* alleles. Strain YPS1009’s *SUC* gene on chromosome VII is colored green at positions where it is identical to strain 273614’s subtelomeric *SUC* allele, purple at positions where it is identical to strain BY’s *SUC2* allele, and orange at positions where it differs from both. **E**. Scatterplots show the growth on sucrose of segregants of the cross between strains 273614 and YJM981, partitioned by *SUC* genotype at chromosomes IV and IX, as shown in Fig. 3. **F**. Growth of segregants between strains 273614 and YJM981 on raffinose, as in panel E. Diagrams of the chemical structures of raffinose and sucrose are shown for reference. Mean phenotypes are demarcated with magenta lines. (NS = not significant, * p < 0.05, one-tailed Student’s *t*-test, with the alternative hypothesis that the extra *SUC* copy improves growth on the tested sugar.)

We also observe four alleles of a *SUC* gene that appears to have been formed through recombination between *SUC2* and the predominant homology group of subtelomeric *SUC* genes (Fig. 5B, D). There are three copies of this intermediate *SUC* gene in strain YPS1009 and one in strain 273614. Interestingly, this *SUC* gene is found both at subtelomeric loci and at the *SUC2* locus, whereas we do not observe the most common subtelomeric *SUC* allele at the *SUC2* locus or the predominant *SUC2* allele at a subtelomere. Nonetheless, the existence of this intermediate *SUC* gene at both loci reinforces that recombination between the subtelomeres and the *SUC2* locus is ongoing.

Previous studies have found that the subtelomeric *SUC* genes are neither silent (Bozdag and Greig 2014) nor nonfunctional, as they can allow strains lacking *SUC2* to grow on sucrose (Carlson and Botstein 1983). Indeed, subtelomeric *SUC* genes can create QTLs for sucrose growth when *SUC2* is nonfunctional. Strain YJM981 has a natural frameshift loss-of-function mutation in *SUC2*, while strain 273614 has both a functional *SUC2* and a subtelomeric *SUC* on chromosome IV. In the 273614 × YJM981 QTL mapping, segregants with YJM981’s *SUC2* frameshift allele grew substantially better on sucrose if they inherited the chromosome IV subtelomeric *SUC* from strain 273614 than if they inherited the *SUC*-less YJM981 chromosome IV (Fig. 5E). In contrast, we noticed that segregants that inherited the functional *SUC2* allele from strain 273614 had no detectable boost to their sucrose growth if they additionally inherited the subtelomeric *SUC* on chromosome IV. This was perplexing, as it suggested that although subtelomeric *SUC*s have maintained invertase function, growth on sucrose may not select for subtelomeric *SUC* function for the majority of the yeast population carrying functional *SUC2*.

Invertase also digests other sugars, so we conjectured that metabolism of other sugars could have selected for the additional invertase activity provided by telomeric *SUC* genes. Raffinose is a trisaccharide composed of galactose, glucose, and fructose. Metabolism of raffinose in yeast involves hydrolysis by invertase to produce fructose and melibiose, a disaccharide of glucose and galactose. Invertase levels are more likely to be limiting for growth in raffinose than sucrose, as invertase has lower activity on raffinose (Gascon et al. 1968; Sainz-Polo et al. 2013). Furthermore, more raffinose would need to be hydrolyzed to produce the same amount of energy, as most yeast strains can only metabolize the resulting fructose and not the melibiose, whereas both the glucose and the fructose produced by sucrose hydrolysis can be consumed.

Indeed, in contrast to sucrose, segregants with functional *SUC2* in the 273614 × YJM981 cross grew significantly better on raffinose if they additionally inherited the subtelomeric *SUC* gene (Fig. 5F). Thus, the extra invertase activity provided by subtelomeric *SUC* genes may be particularly beneficial for growth in complex sugar environments.

## Discussion

There has been increasing interest in describing species genetically not only through a single reference genome, but with a population-wide pangenome that accounts for genes variably present in individuals. We used long-read sequencing to generate highly complete genome assemblies for 16 strains of *S. cerevisiae* and annotate pangenomic variation. To ascertain the effects of the uncovered genetic variation, we combined our long-read-derived genome sequences with linkage mapping data and identified QTLs caused by variable pangenomic genes. By leveraging linkage mapping data, we were also able to validate the long-read genome sequences, confirming a translocation and the placement of variable pangenomic genes through linkage to nearby genetic markers. Thus, combining linkage mapping and long-read sequencing strengthens our understanding of both the sequenced genomes and mapped QTLs.

In budding yeast, variable pangenomic genes are mostly found at subtelomeric locations (Yue et al. 2017). This could be related to yeast subtelomeres recombining between chromosomes. Interchromosomal recombination can lead to addition or loss of genes following cell division (Louis et al. 1994). The genes we examined here, *MALR, SGE1*, and *SUC*, are all subtelomeric. Consistent with the subtelomeres experiencing interchromosomal recombination, copies of all three genes were found at the end of more than one chromosome. Surveying more strains than the 16 studied here would likely uncover additional genomic localizations of these genes. In some cases, we found identical gene copies on different chromosomes, suggesting their recombination events occurred recently. For *SGE1*, three distinct alleles seen on chromosome XVI are also seen on other chromosomes, implying three independent recombination events, further underscoring the frequency of interchromosomal recombination. Interestingly, subtelomeric interchromosomal recombination has also been observed in humans (Guarracino et al. 2022), and thus could contribute to human trait variation.

Increased gene dosage of limiting gene products can increase fitness, and we observed that increasing from one functional gene to two improved fitness through *MALR* for maltose, *SGE1* for paraquat, and *SUC* for raffinose. The variable presence of highly identical sequences on different chromosomes is particularly challenging to detect using reference genomes and short-read genotyping. Correctly determining the genomic locations of variable genes is especially important for resolving the genetic basis of mapped QTLs, which requires precise knowledge of the sequence differences in the QTL. QTLs were caused by genes in locations absent from the reference genome for all three traits examined here.

We predict that variable pangenomic genes have many more effects on yeast phenotypic diversity than the examples we described here. The variable component of the yeast pangenome comprises thousands of genes (Peter et al. 2018), and each of those genes may contribute to very particular traits, such as metabolism of specific nutrients or growth in specific environments. For instance, several homologs of *MALR* were widespread in our 16 genomes but did not support growth on maltose, raising the possibility that they contribute to fitness in a different sugar environment. With the variable pangenomic content annotated here for a large, genotyped mapping population, studying a wide variety of phenotypes in the panel’s segregants could reveal the phenotypic effects of many other pangenomic genes in these genomes.

The biparental cross structure of the study population meant that generating high-quality genome assemblies for the parental strains, rather than all the segregants, was sufficient for resolving QTLs. In addition to biparental crosses, this strategy can be utilized for linkage mapping studies on other groups of related individuals, such as family studies in humans. As the cost and analytical constraints of generating long-read genome assemblies continues to decline (Jarvis et al. 2022), it will become possible to generate high-confidence de novo whole genome sequences of every individual in a mapping study, including genome-wide association studies of unrelated humans. We expect that assembly of highly complete genomes using long-read sequencing will be vital for us to understand the genetic sources of trait variation.

## Acknowledgements

We thank Arang Rhie, Sergey Koren, Adam Phillippy, Bill Pavan, David Bodine, Frank Albert, and all members of the Sadhu lab for helpful discussion. We thank Trey Sato, Leonid Kruglyak, and Fred Cross for strains and plasmids. This work was supported by the Intramural Research Program of the National Human Genome Research Institute, National Institutes of Health (1ZIAHG200401, 1ZIBHG000196). J.S.B. was supported by the Howard Hughes Medical Institute.

## Author contributions

M.J.S. conceived and supervised the project. J.S.B. and M.J.S. provided resources. I.A., C.A.W., M.C., and M.J.S performed investigation and formal analysis. M.P. and the NISC Comparative Sequencing Program sequenced and assembled yeast genomes. M.J.S. wrote the original draft; all authors reviewed and edited the manuscript and approved the final version.

## Materials and Methods

### Genome sequencing and assembly

DNA was isolated from the 16 parental strains using the Masterpure Yeast DNA Purification Kit (Lucigen, Middleton, WI, USA). The Pacific Biosciences protocol “Preparing HiFi SMRTbell® Libraries using SMRTbell Express Template Prep Kit 2.0” was used to create libraries from 30 micrograms of DNA. The Megarupter (Diagenode, Denville, NJ, USA) was used for shearing and Blue Pippin (Sage Science, Beverly, MA, USA) was used for size-selection for fragments greater than 13 kb. A subset of the libraries was constructed using barcoded adapters to allow pooling before size-selection.

Each library was run on one SMRTCell (version 8M) using version 2.0 sequencing reagents. Sequencing was performed on a Sequel II sequencer (Pacific Biosciences, Menlo Park, CA, USA) running instrument control software version 9.0.0.9223 and with a movie collection time of 30 hours per SMRTCell.

For individual sample libraries, circular consensus sequence (CCS) reads, also known as HiFi reads, were generated from the initial subread data using the pb_ccs workflow (ccs version 4.2.0) within PacBio SMRTLink version 9.0.0.92188. For the pooled libraries, demultiplexed CCS reads were generated from the initial subread data using the pb_demux_ccs workflow (lima version 1.11.0) within PacBio SMRTLink.

CCS reads from each sample were assembled using CANU version 2.2 (Koren et al. 2017) (parameters `genomeSize=12mb-pacbio-hifi`). In cases where read coverage was greater than 50x, the CCS reads were downsampled to 50x coverage using seqtk v1.2 before assembly (https://github.com/lh3/seqtk). Redundant contigs were identified from each initial draft assembly using purge_dups v1.0.1 (Guan et al. 2020). The mitochondrial contig sequences were visualized and manually circularized using Gepard v1.30 (Krumsiek et al. 2007). Further refinement of the contig sequences involved comparing the draft contigs against the Saccharomyces cerevisiae S288C reference genome (ASM205763v1) using BLASTn (v2.2.24) to identify contigs representing telomeric ends that were not incorporated into the corresponding chromosome and overlapping chromosome contigs, particularly the two contigs representing chromosome XII that were split at the rDNA cluster. Overlapping contigs were visualized using Gepard to identify the overlapping regions and manually joined using exported trimmed sequences. The structure of the rDNA cluster in the joined chromosome XII contigs represents a collapsed version of this extended repeat region.

### Genome annotation

Genes were annotated in the strain fasta files using a combined strategy of lifting over gene annotations from the reference *S. cerevisiae* genome and annotating ORFs de novo. For lifting over reference annotations, we used Liftoff v.1.6.1 (Shumate and Salzberg 2021) with the parameters -p 4 -infer_transcripts -copies -s .3 -sc .3. The reference annotations were downloaded from https://downloads.yeastgenome.org/latest/saccharomyces_cerevisiae.gff.gz on December 1, 2021.

For annotating ORFs de novo, we used ORFfinder v.0.4.3 with the parameters -outfmt=0 -ml 210. Using custom R scripts (R v.4.1.2), ORFs found by ORFfinder were filtered to exclude ORFs that overlapped other genomic features, namely genes, transposable elements, centromeres, tRNAs, subtelomeric X and Y′ elements, and autonomously replicating sequences. The sites of these features in each genome were determined by lifting them over from the reference genome using Liftoff, after classifying their feature types in the reference annotations as “gene.” We additionally filtered for a minimum ORF length of 100 codons, and, as ORFfinder includes ORFs with non-canonical start codons, we filtered for ORFs that began with an ATG. Remaining ORFs were combined into .gff files outputted by Liftoff.

In addition to being used in filtering ORFfinder ORFs, the lifting over of additional feature types was used for determining which contigs ended in subtelomeric X and Y′ elements in Figure 1A, by checking for their presence in the first 30kb or after the first 200 kb of each contig.

### Validation of assemblies using genetic maps

For a given cross and chromosome, we used MUMmer (v4.0.0) (Marcais et al. 2018) to generate a chromosome-to-chromosome aligned reference sequence between the chromosomes of interest from the two parent genome sequences, and call SNPs. Using custom scripts, we only included SNPs that were at least 20 nucleotides away from an alignment mismatch (including indels), and replaced invariant or poorly-aligned regions with Ns to yield a final reference sequence. Illumina short reads of the segregants, obtained from the Sequence Read Archive (study SRP201965), were aligned with bwa mem (v0.7.17) (Li and Durbin 2009) using default parameters and SNPs were called with bcftools (v.1.16) (Li 2011) using default parameters. To reduce noise from SNP calling, we called genotypes for non-overlapping 1000 bp windows, assigning parentage if at least 90% of SNPs within a window agreed on parentage (otherwise assigned N). Linkage disequilibrium and association mapping were done with plink (v1.9) (Chang et al. 2015). For all tests, we converted genotype calls into .ped and .map files using custom R and Python scripts.

For testing whether particular gene sequences of interest were assembled onto the correct chromosome, we performed linkage mapping in segregants of biparental crosses predicted to be heterozygous for a sequence of interest. We counted Illumina reads for each segregant in a cross that matched the gene sequence. Because of the high similarity of these gene sequences to other homologs, we employed strict bwa mapping parameters to allow only precise matches: mismatch penalty of 40 (-B, default=4); gap open penalty of 60 (-O, default=6); gap extension penalty of 10 (-E, default=1); and end clipping penalty of 100 (-L, default=5). Linkage mapping for these sequences of interest incorporated the number of reads mapped to the diagnostic reference sequence as a quantitative phenotype (reads mapped per million, RPM).

### QTL and segregant analysis

All data used to generate LOD plots for maltose and paraquat was generated by Bloom et al. In brief, Bloom et al. calculated segregant phenotypes as the size of segregant colonies after 48 hours of growth on agar plates presenting a growth condition of interest, then normalized against growth on YPD plates. Normalized phenotypes were scaled to have mean of 0 and variance of 1 for a given cross and condition. Bloom et al. determined segregant genotypes from 100-bp or 150-bp Illumina reads using the GATK haplotype caller (McKenna et al. 2010), then imputed missing genotypes and structural variant genotypes from the parental genotypes using a hidden Markov model. Phenotype and genotype data were downloaded from https://www.dropbox.com/sh/jqm7a11zz9laytd/AABaE0EfQxLH6ounPhJ7yYWya. We calculated LOD scores at each biallelic marker as:

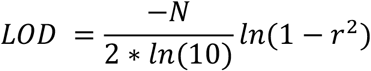

where *r* is the Pearson correlation between the phenotypes and genotypes of the segregants in the cross, and *N* is the number of segregants; see page 454 of Lynch and Walsh (Lynch and Walsh). Strain CLIB219 has a loss-of-function mutation in *ADE2* that dominates the QTL mapping of nearly all traits involving CLIB219; as such, we filtered out 119 segregants with the *ade2-* genotype from the CBS2888 × CLIB219 cross and 123 segregants from the CLIB219 × M22 cross, then performed all phenotypic analyses on these crosses using the remaining 824 and 820 segregants, respectively.

For scatterplots of segregant phenotypes, we partitioned segregants by genotypes called by Bloom et al. of SNPs close to the QTLs being examined. SNPs were chosen because we felt the Illumina-sequencing-based genotype-calling would have higher confidence calls for SNPs. Chosen SNPs were also checked for segregation ratios in the segregants close to the expected 50:50, and for well-behaved linkage disequilibrium patterns with neighboring variants.

### Plasmid construction

Strains, plasmids, and oligonucleotides are listed in Supplemental Tables 2, 3, and 4, respectively. The chromosome VII *MALR* from strain YJM145 (MSY28), referred to as *MALR*_YJM145_, was cloned by Gibson assembly (Gibson et al. 2009) onto pRS418 (MSp183, a gift of Fred Cross), a CEN plasmid with a *NatMX* selectable marker. First, *MALR*_YJM145_ was PCR-amplified from YJM145 genomic DNA isolated using a DNeasy blood and tissue kit (Qiagen, Valencia, CA, USA). The PCR used primers oMS90/111, which spanned from 827 bp upstream to 690 bp downstream of the *MALR*_YJM145_ ORF, and the Herculase II Fusion DNA polymerase (Agilent, Santa Clara, CA, USA). Meanwhile, pRS418 was digested using ApaI and EagI-HF (New England Biolabs, Ipswich, MA, USA). The PCR product was PCR-purified and the plasmid digest was gel-extracted (Qiagen), after which the two were assembled together using Gibson assembly (New England Biolabs) to produce plasmid MSp198. The sequence of *MALR*_YJM145_ on MSp198 was confirmed with Sanger sequencing (Eurofins Genomics, Louisville, KY, USA).

*SGE1* alleles from strains BY, YJM145, YPS1009, and I14 (MSY25, 28, 32, and 33, respectively) were cloned by Gibson assembly onto pRS415 (MSp8), a CEN plasmid with a *LEU2* selectable marker. PCRs of the alleles were generated with oligonucleotides oMS100/101 and spanned from 467 bp upstream to 357 bp downstream of the *SGE1* ORF. pRS415 was digested with EagI-HF and HindIII-HF (New England Biolabs), and cloning was performed as for *MALR*_YJM145_ above. The *SGE1* allele from strain 273614 was generated by introducing strain 273614’s L449M substitution into the plasmid carrying *SGE1*_14_ (MSp191). MSp191 was amplified with primers oMS128/129 using Herculase II Fusion DNA polymerase, after which the resulting PCR product was treated with DpnI (New England Biolabs) and PCR-purified. Gibson assembly was performed using the PCR product and oligonucleotide oMS127 to produce the plasmid carrying the *SGE1*-L449M substitution in strain 273614.

### Maltose growth assay

The maltose growth assay was performed on strains BY, YJM987, YPS1009, Y10, PW5, and CLIB219 (MSY25, 29, 31, 32, 35, and 39, respectively) transformed with either empty vector (MSp183) or the plasmid carrying *MALR*_YJM145_ (MSp198) using standard lithium acetate transformation procedures (Becker and Lundblad 2001). As strains MSY32 and MSY39 are annotated as carrying plasmids with *NatMX* markers, prior to transformation they were plated without selection for the plasmid, from which resulting nourseothricin-sensitive colonies were used for transformations (GoldBio, Saint Louis, MO, USA). Five distinct colonies from each transformation plate were used as experimental replicates, except for YJM978 + empty vector, which had 4 colonies total.

Single colonies of strains were grown for two days without shaking at 30C in a 96-well deep-well plate (Thermo Fisher Scientific, Waltham, MA, USA; catalog no. 12566612) in 500 μL YPD media with added adenine sulfate (20 mg/L) and nourseothricin (100 μg/mL). The position of each strain on the plate was randomly assigned, and the first and last columns were filled with water, not cultures. One microliter of saturated yeast cultures were transferred into the equivalent position of three 96-well clear flat-bottom plates (Corning, Corning, NY, USA; catalog no. 3370) filled with 198 μL YNB media without ammonium sulfate (BD, Franklin Lakes, NJ, USA; catalog no. 233520) with added monosodium glutamate (Sigma-Aldrich, Burlington, MA, USA; catalog no. G1626), adenine, nourseothricin, as well as 20 g/L maltose, 20 g/L glucose, or no added sugar. Absorbance (OD_600_) of each well of each plate was measured in parallel on a SPECTROstar Omega microplate reader equipped with a plate stacker (BMG Labtech, Ortenberg, Germany) over a period of 93 hours at ambient temperature (∼25 C) with 30 s of shaking at 600 RPM prior to each plate reading. Reads were taken approximately 14 minutes apart.

For each sample, the growth phenotype was calculated as the ratio of the area under the growth curve in maltose media (AUC_maltose_) to the area under the growth curve in glucose media (AUC_glucose_), i.e. AUC_maltose_/AUC_glucose_. AUC values were calculated by summing the curve’s OD_600_ values and then subtracting the summed OD_600_ values for the same sample’s no-sugar growth curve as a blank normalization. For figures, OD_600_ values were normalized to OD_600_ values in media without sugar then plotted using the smooth.spline function in R with the parameter lambda=0.0001, though AUC calculations were performed on unsmoothed growth curves. Statistical analysis was performed using one-way ANOVA followed by Tukey’s post-hoc HSD test.

### Paraquat growth assay

The paraquat growth assay were performed on strain *sge1Δ* from the BY MATa knockout collection (Transomic, Huntsville, AL, USA) transformed with plasmids carrying different alleles of *SGE1* (MSp189-192 and 200), as well as empty vector (MSp8). Ten distinct colonies from each transformation plate were seeded as experimental replicates into a 96-well deep-well plate, with the outer rows and columns filled with water instead of cultures. Positions of replicates on the plate were evenly distributed across rows and columns, rather than randomized, to avoid chance association of a strain with a particular row or column of the plate. For instance, the replicates of the strain with *SGE1*_273614_ were in wells G2, F3, E4, D5, etc. The media was YNB plus glucose with CSM minus uracil and leucine (Sunrise Science, Knoxville, TN) with uracil (20 mg/L) added back.

We had previously observed that paraquat had an inconsistent effect on yeast growth in the plate reader, with lower toxicity in rows towards the bottom of the plate. We used a plate arrangement in which all growth data was generated using the top row of the culture plate. After two days of growth, 2 μL of the precultures were seeded into seven plates to go onto the plate reader: six plates with 198 μL YPD containing 3.5 mM paraquat, and one plate with YPD without paraquat. Note that a higher concentration of paraquat was used here for the liquid media growth assays than was used by Bloom et al. to measure plate-based paraquat growth, 3.5 mM instead of 0.75 mM, because 0.75 mM paraquat had little effect on yeast growth in the liquid media assay. The positions of the cultures in the six plates with 3.5 mM paraquat were rotated by row: plate 1 had cultures seeded into equivalent positions from the preculture plate, then plate 2 had the bottom preculture row seeded into its top culture row with the other culture rows seeded one row down, etc. Thus, across the six paraquat plates, all samples were in the top culture row once. Culture density readings were taken approximately 32 minutes apart for 113 hours. Only the top row of each paraquat plate was used in downstream analyses.

For each sample, the growth phenotype was calculated as the ratio of the area under the growth curve in paraquat media (AUC_paraquat_) to the area under the growth curve in media without paraquat (AUC_no paraquat_), i.e. AUC_paraquat_/AUC_no paraquat_. AUC values were calculated by summing the OD_600_ values for the first 96 hours after subtracting from each value the OD_600_ reading at *t* = 0 as a blank normalization.

## Data deposition

PacBio-based genome assemblies of the strains in our 16-isolate panel are deposited at the NCBI genome database with accession numbers SAMN23081757 through SAMN23081772 (see Supplemental Table 1). Code used to generate and process our data is available at an up-to-date repository hosted at https://github.com/cory-weller/long-read-yeast-genomes, and will be archived in a permanent, independent repository at Zenodo prior to publication.

