## Supplemental Figures for "Long-read genomes reveal pangenomic variation underlying yeast phenotypic diversity"

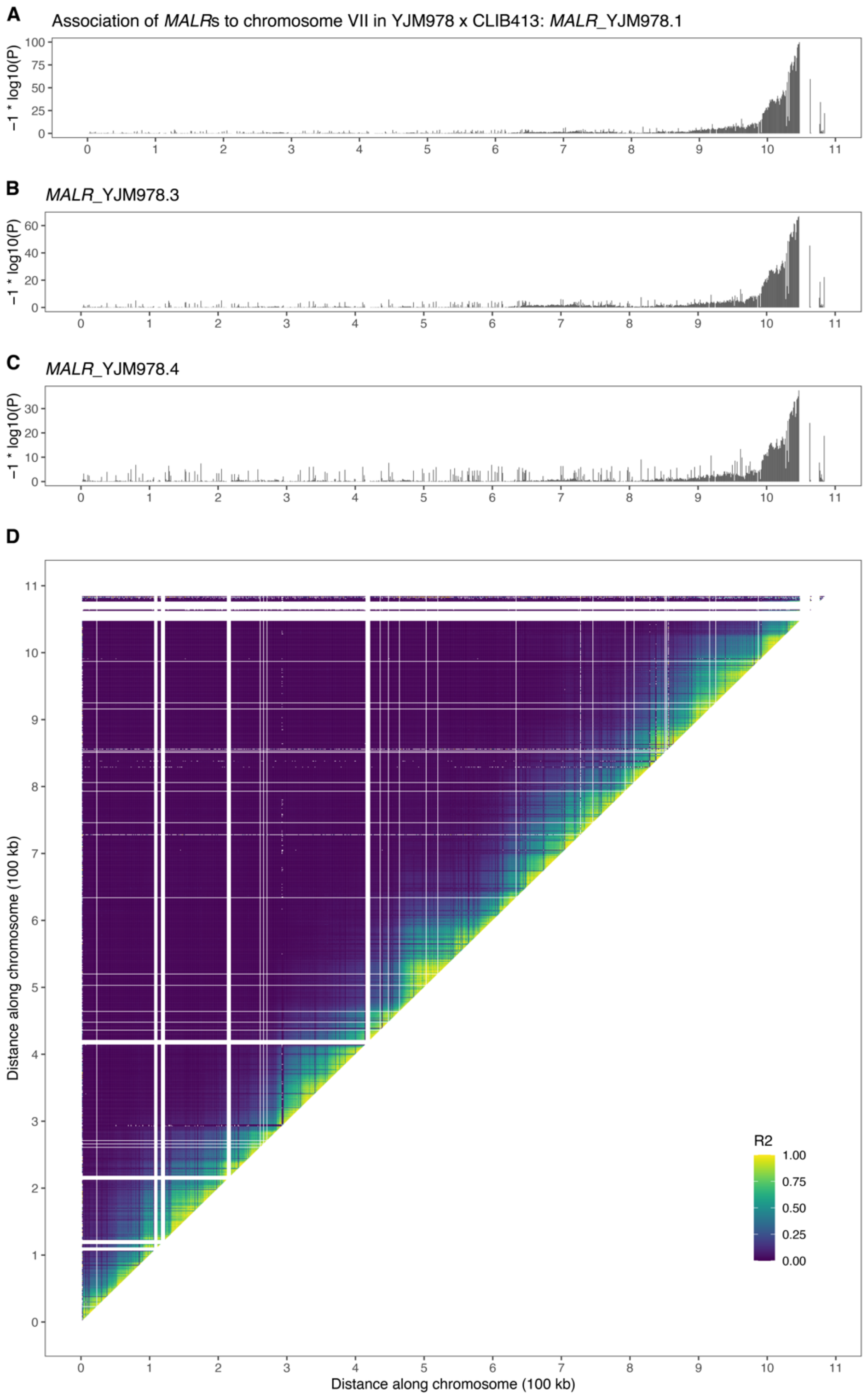


**Figure S1. Validation of assembly of YJM978 *MALR*s onto chromosome VII-R through linkage mapping in YJM978 × CLIB413.**

**A-C.** Chromosome VII linkage mapping of $MALR_{YJM978.1}$, $MALR_{YJM978.3}$, and $MALR_{YJM978.4}$, representing *MAL13*, *MAL64*, and *MAL63*, respectively. For each segregant in the YJM978 × CLIB413 cross, the presence of Illumina short reads matching the query gene sequence was determined using bwa, and quantified as reads per million. This value was treated as a quantitative trait and mapped in the YJM978 × CLIB413 segregant panel using SNPs on chromosome VII segregating between YJM978 and CLIB413. Association values are plotted against chromosome VII coordinates. Note that $MALR_{YJM978.2}$ (representing *MAL73*) was not tested, as *MAL73* is present on the right arm of both the YJM978 and CLIB413 chromosome VII assemblies and thus is predicted to be inherited in 100% of segregants, which blocks the use of linkage to confirm its chromosomal location.

**D.** Linkage disequilibrium plot of segregating genotypes on chromosome VII in YJM978 × CLIB413. There is high correlation along the entire diagonal with no completely disjoint linkage blocks. This indicates that the chromosome VII assemblies in YJM978 and CLIB413 did not incorrectly concatenate a different chromosome's sequence onto the end of chromosome VII, which could lead to misannotation of *MALR*s onto chromosome VII. White segments denote missing data due to the lack of suitable SNPs for genotyping segregants in that window.

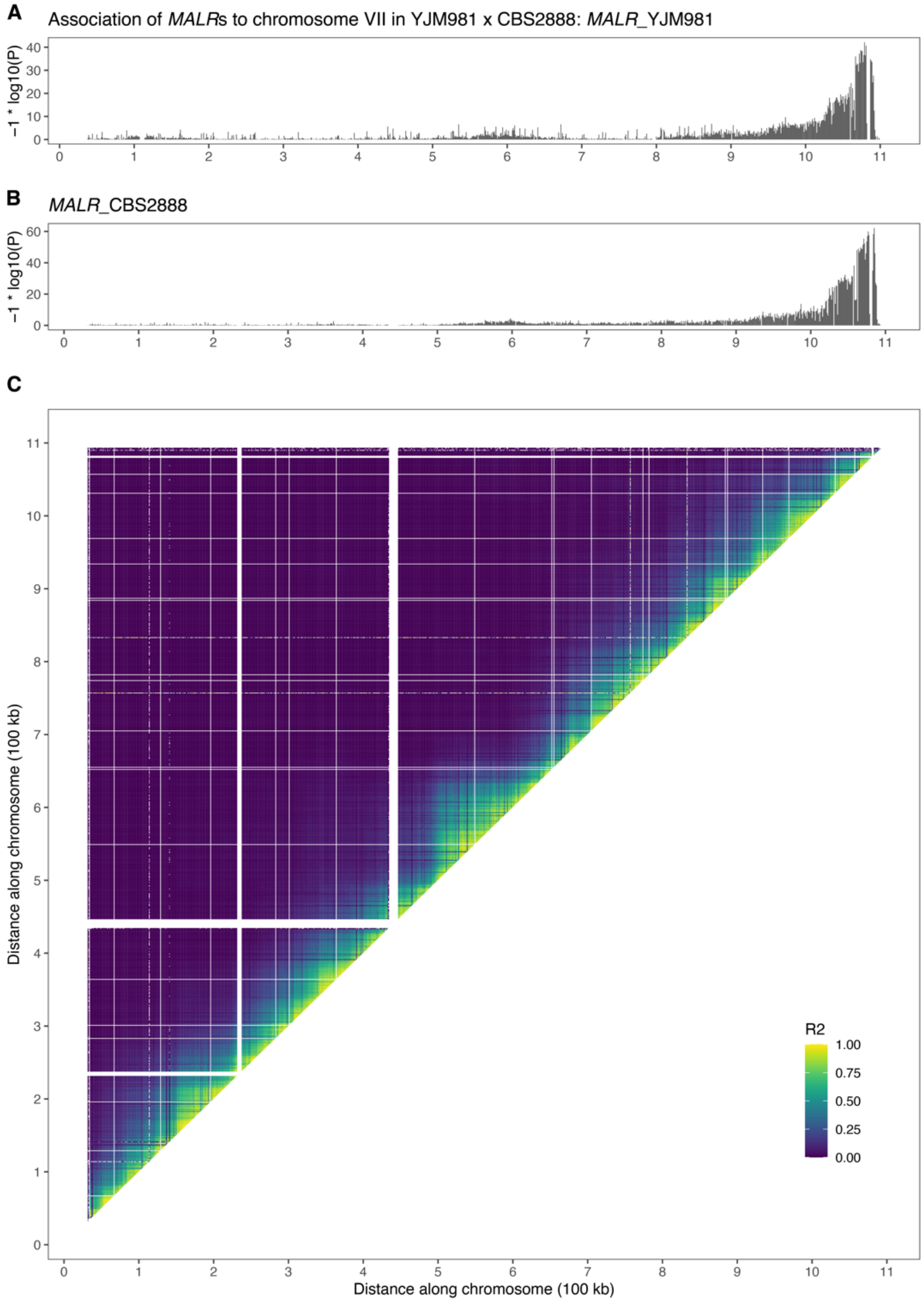


**Figure S2. Validation of assembly of *MALR*$_{YJM981}$ and *MALR*$_{CBS2888}$ onto chromosome VII-R through linkage mapping in YJM981 × CBS2888.**

**A-B.** Chromosome VII linkage mapping of *MALR*$_{YJM981}$ and *MALR*$_{CBS2888}$, representing *MAL23* and *MAL73*, respectively. Association is shown as in Supplemental Fig. 1A-C.

**C.** Linkage disequilibrium plot of segregating genotypes on chromosome VII in YJM981 × CBS2888. Linkage disequilibrium is shown as in Supplemental Fig. 1D.

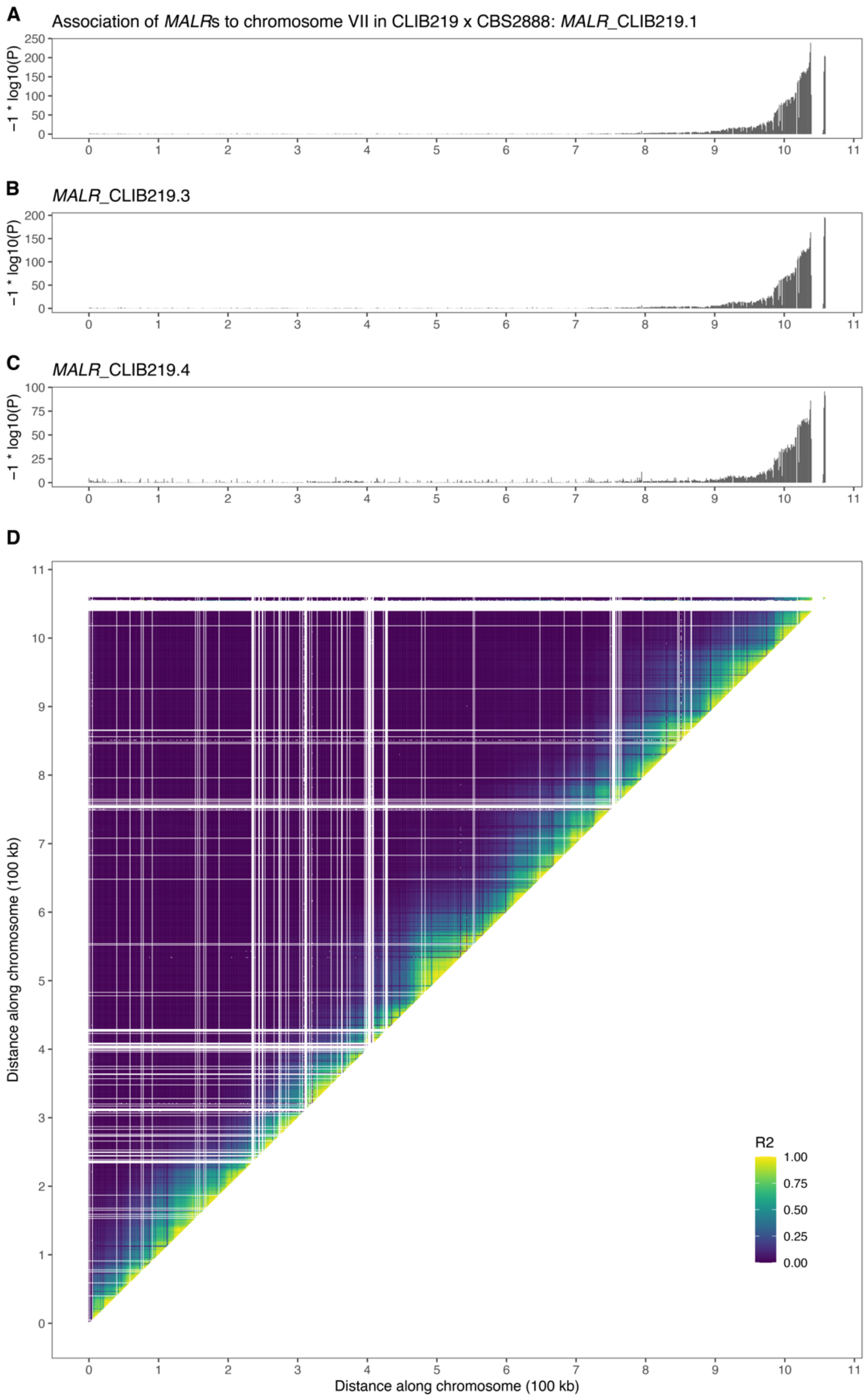


**Figure S3. Validation of assembly of CLIB219 *MALR*s onto chromosome VII-R through linkage mapping in CLIB219 × CBS2888.**

**A-C.** Chromosome VII linkage mapping of *MALR*$_{\text{CLIB219.1}}$, *MALR*$_{\text{CLIB219.3}}$, and *MALR*$_{\text{CLIB219.4}}$, representing *MAL13*, *MAL64*, and *MAL43*, respectively. Association is shown as in Supplemental Fig. 1A-C. As in Supplemental Fig. 1, *MALR*$_{\text{CLIB219.2}}$ (representing *MAL73*) was not tested, as *MAL73* is present on the right arm of both the CLIB219 and CBS2888 chromosome VII assemblies.

**D.** Linkage disequilibrium plot of segregating genotypes on chromosome VII in CLIB219 × CBS2888. Linkage disequilibrium is shown as in Supplemental Fig. 1D.

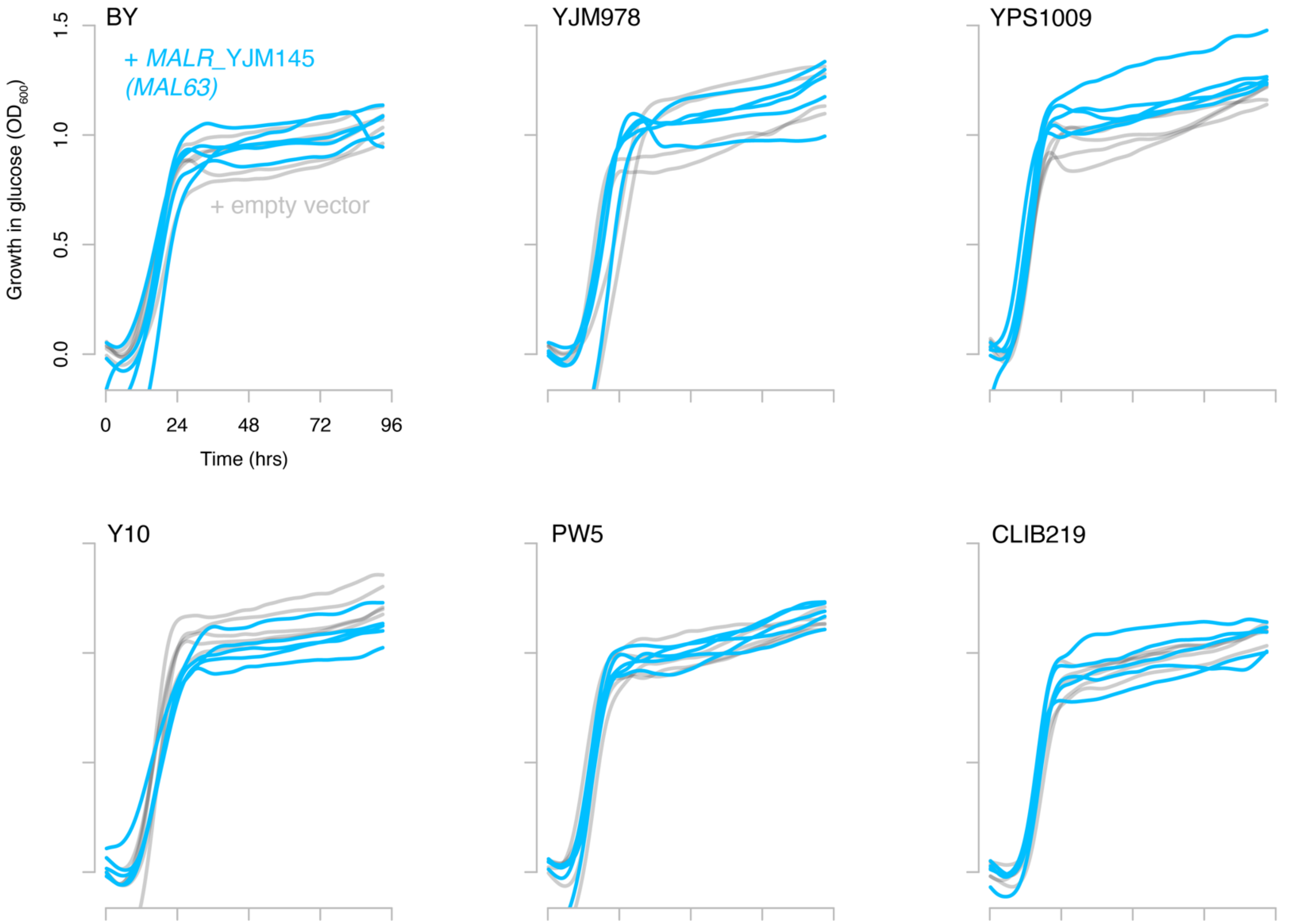


**Figure S4. Growth in glucose is unaffected by *MALR* content.**
Growth over time in glucose for strains BY, YJM978, YPS1009, Y10, PW5, and CLIB219 transformed with a plasmid expressing *MALR*$_{YJM145}$ (blue) or empty vector (grey). n = 5 biological replicates, except for YJM978 + empty vector, which had 4 biological replicates. Background absorbance was subtracted, calculated from the ODs of the strains grown without sugar.

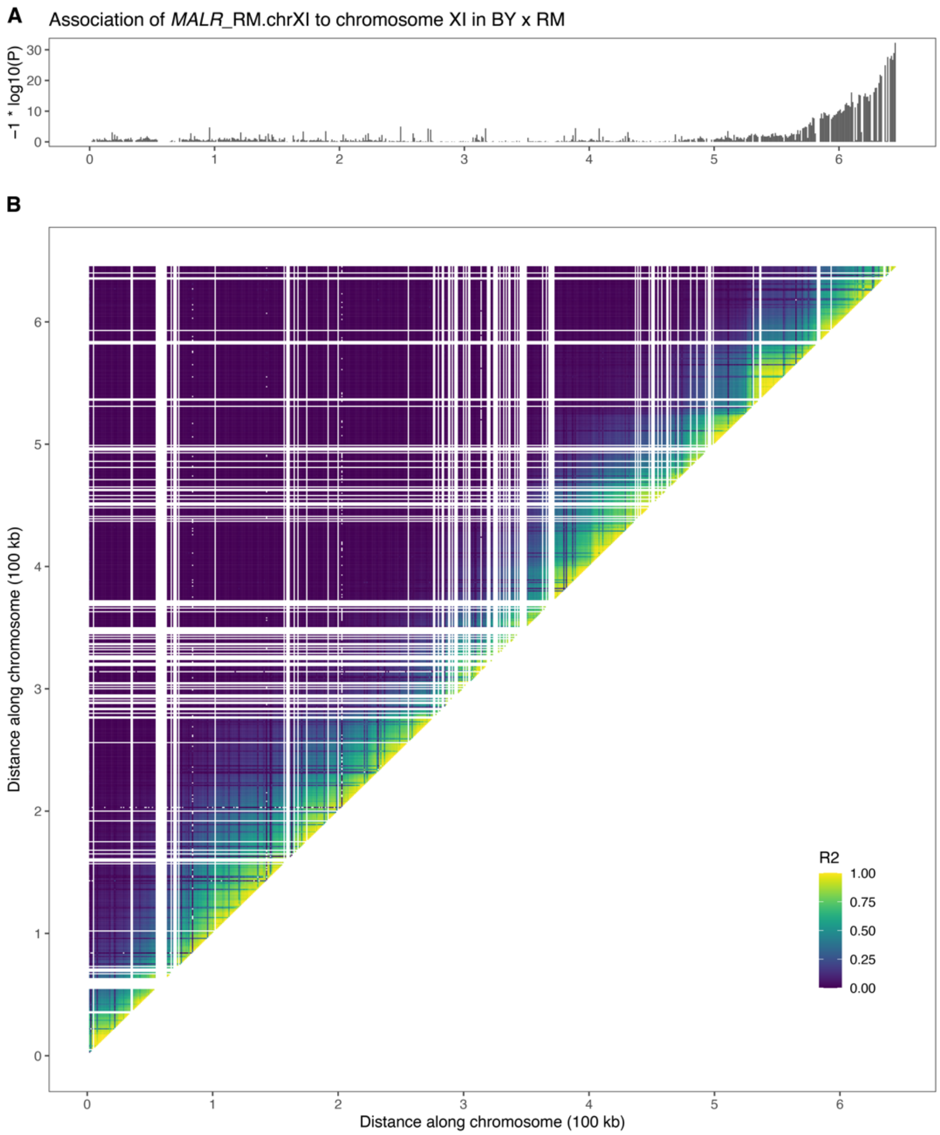


**Figure S5. Validation of assembly of *MALR*$_{\mathbf{RM.chrXI}}$ onto chromosome XI-R through linkage mapping in BY × RM.**

**A.** Chromosome XI linkage mapping of *MALR*$_{\mathrm{RM.chrXI}}$. Association is shown as in Supplemental Fig. 1A-C.

**B.** Linkage disequilibrium plot of segregating genotypes on chromosome XI in BY × RM. Linkage disequilibrium is shown as in Supplemental Fig. 1D.

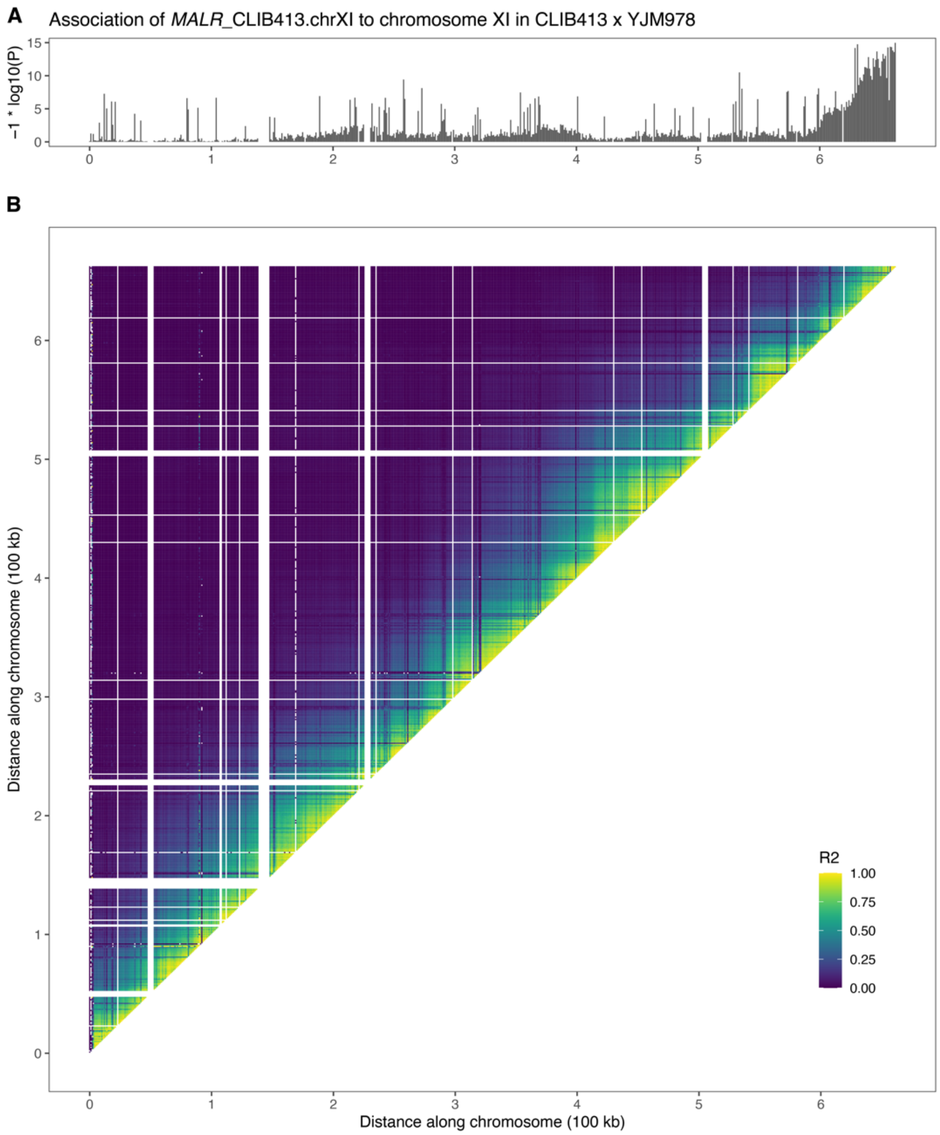


**Figure S6. Validation of assembly of *MALR*$_{\mathbf{CLIB413.chrXI}}$ onto chromosome XI-R through linkage mapping in CLIB413 × YJM978.**

**A.** Chromosome XI linkage mapping of *MALR*$_{\mathrm{CLIB413.chrXI}}$. Association is shown as in Supplemental Fig. 1A-C.

**B.** Linkage disequilibrium plot of segregating genotypes on chromosome XI in CLIB413 × YJM978. Linkage disequilibrium is shown as in Supplemental Fig. 1D.

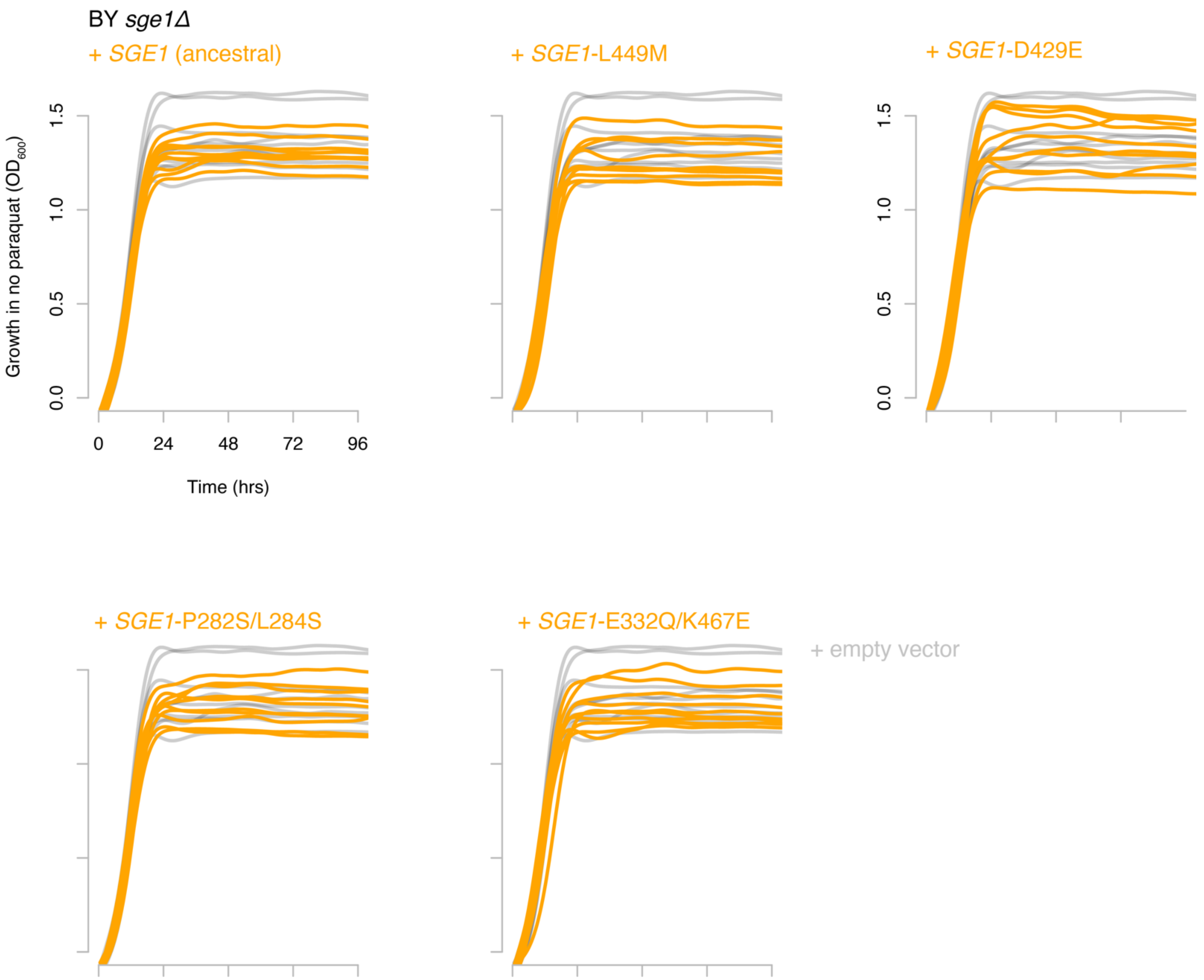


**Figure S7. Expression of *SGE1* alleles does not affect growth in media without paraquat.** Growth over time in liquid YPD media for BY *sge1Δ* transformed with plasmids expressing *SGE1* alleles from strains BY, YJM145, YPS1009, I14, or 273614 (orange). Growth curves are overlaid onto growth curves of *sge1Δ* transformed with empty vector (grey) for comparison. $n$ = 10 biological replicates. Background absorbance at $t$ = 0 was subtracted from all other timepoints for each curve.

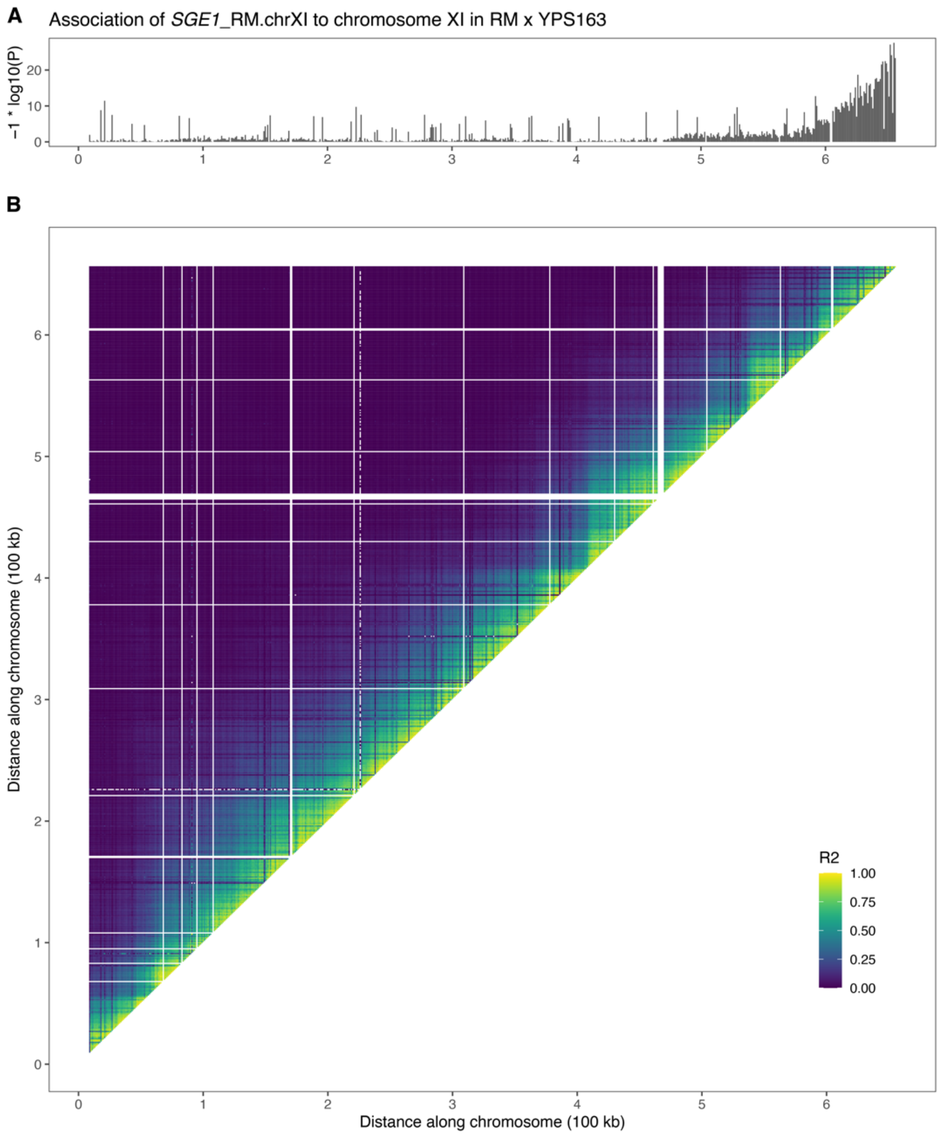


**Figure S8. Validation of assembly of *SGE1*$_{\text{RM.chrXI}}$ onto chromosome XI-R through linkage mapping in RM × YPS163.**
**A.** Chromosome XI linkage mapping of *SGE1*$_{\text{RM.chrXI}}$. Association is shown as in Supplemental Fig. 1A-C.
**B.** Linkage disequilibrium plot of segregating genotypes on chromosome XI in RM × YPS163. Linkage disequilibrium is shown as in Supplemental Fig. 1D.

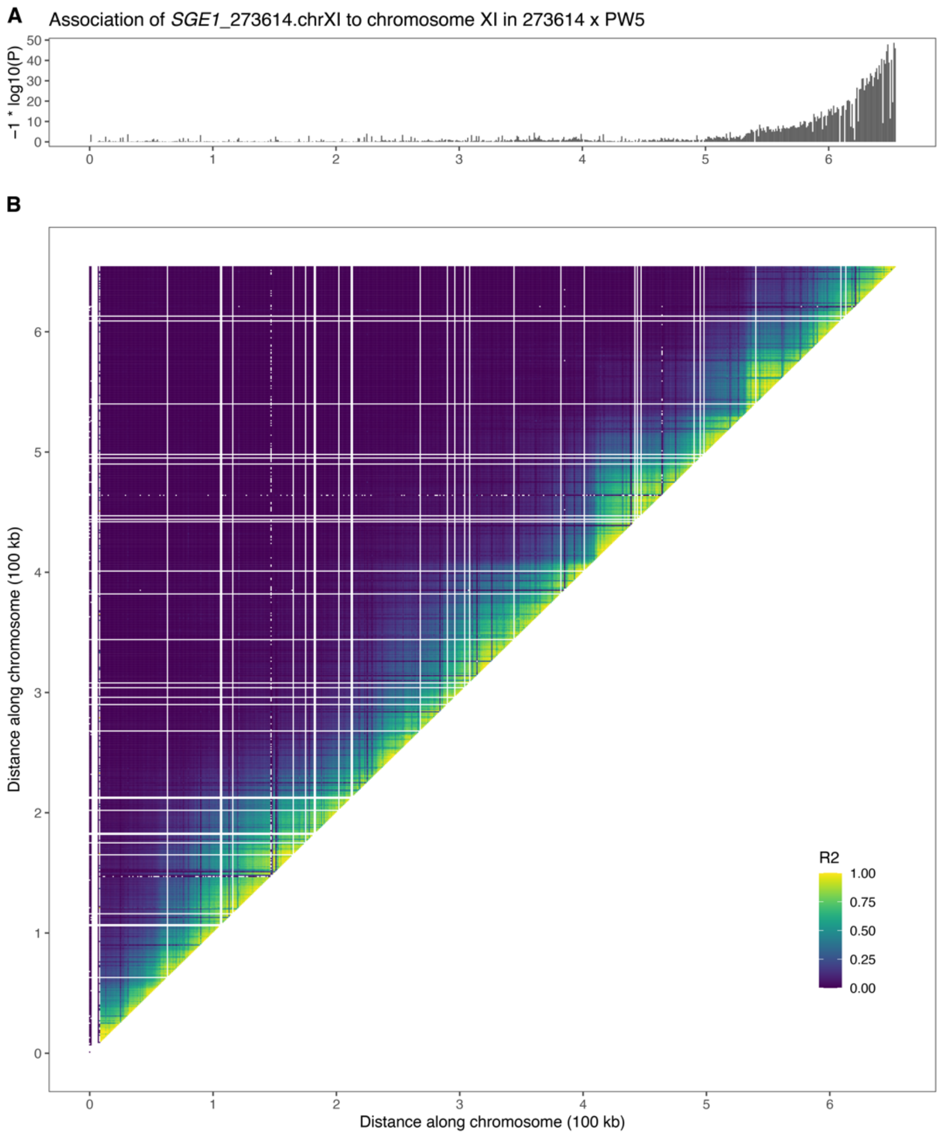


**Figure S9. Validation of assembly of *SGE1*$_{273614.chrXI}$ onto chromosome XI-R through linkage mapping in 273614 × PW5.**

**A.** Chromosome XI linkage mapping of *SGE1*$_{273614.chrXI}$. Association is shown as in Supplemental Fig. 1A-C.

**B.** Linkage disequilibrium plot of segregating genotypes on chromosome XI in 273614 × PW5. Linkage disequilibrium is shown as in Supplemental Fig. 1D.

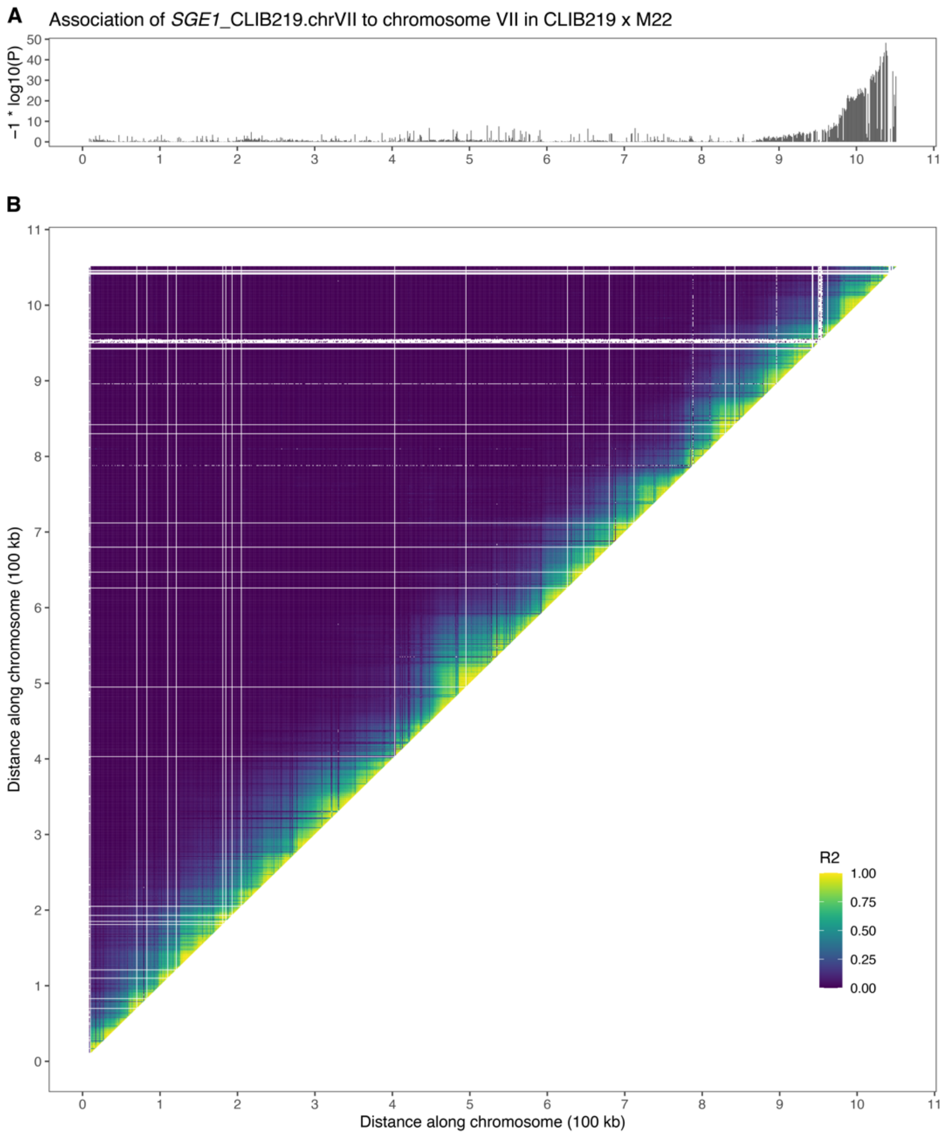


**Figure S10. Validation of assembly of *SGE1*$_{CLIB219.chrVII}$ onto chromosome VII-R through linkage mapping in CLIB219 × M22.**
**A.** Chromosome VII linkage mapping of *SGE1*$_{CLIB219.chrVII}$. Association is shown as in Supplemental Fig. 1A-C.
**B.** Linkage disequilibrium plot of segregating genotypes on chromosome VII in CLIB219 × M22. Linkage disequilibrium is shown as in Supplemental Fig. 1D.

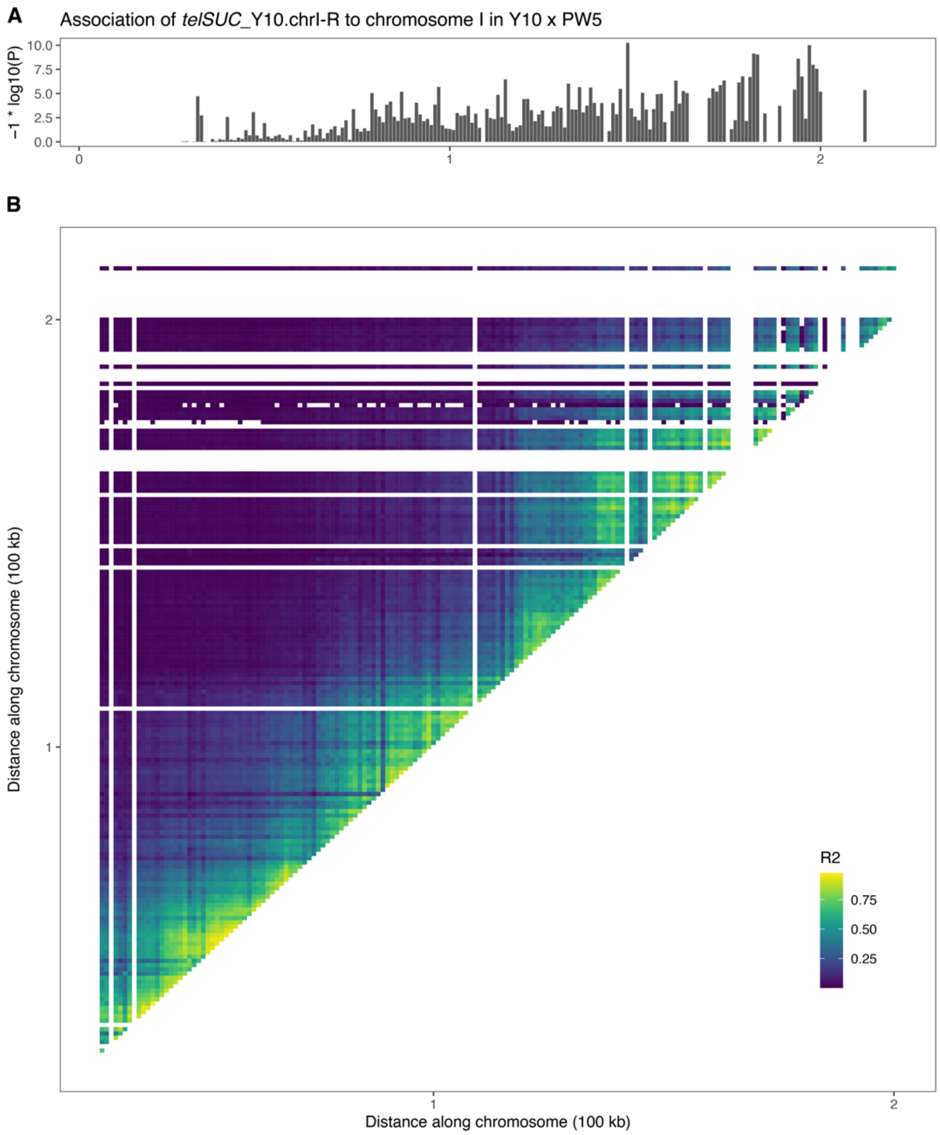


**Figure S11. Validation of assembly of *telSUC*$_{Y10.chrI-R}$ onto chromosome I-R through linkage mapping in Y10 × PW5.**

**A.** Chromosome I linkage mapping of *telSUC*$_{Y10.chrI-R}$. Association is shown as in Supplemental Fig. 1A-C. The linkage is highly significant, but the strength of linkage is lower here than for other analyses due to the presence of identical and near-identical *telSUC* sequences at other Y10 chromosome ends. The linkage signal is also spread over a larger component of the chromosome here than for other analyses due to chromosome I being very short and thus experiencing fewer recombination events.

**B.** Linkage disequilibrium plot of segregating genotypes on chromosome I in Y10 × PW5. Linkage disequilibrium is shown as in Supplemental Fig. 1D.

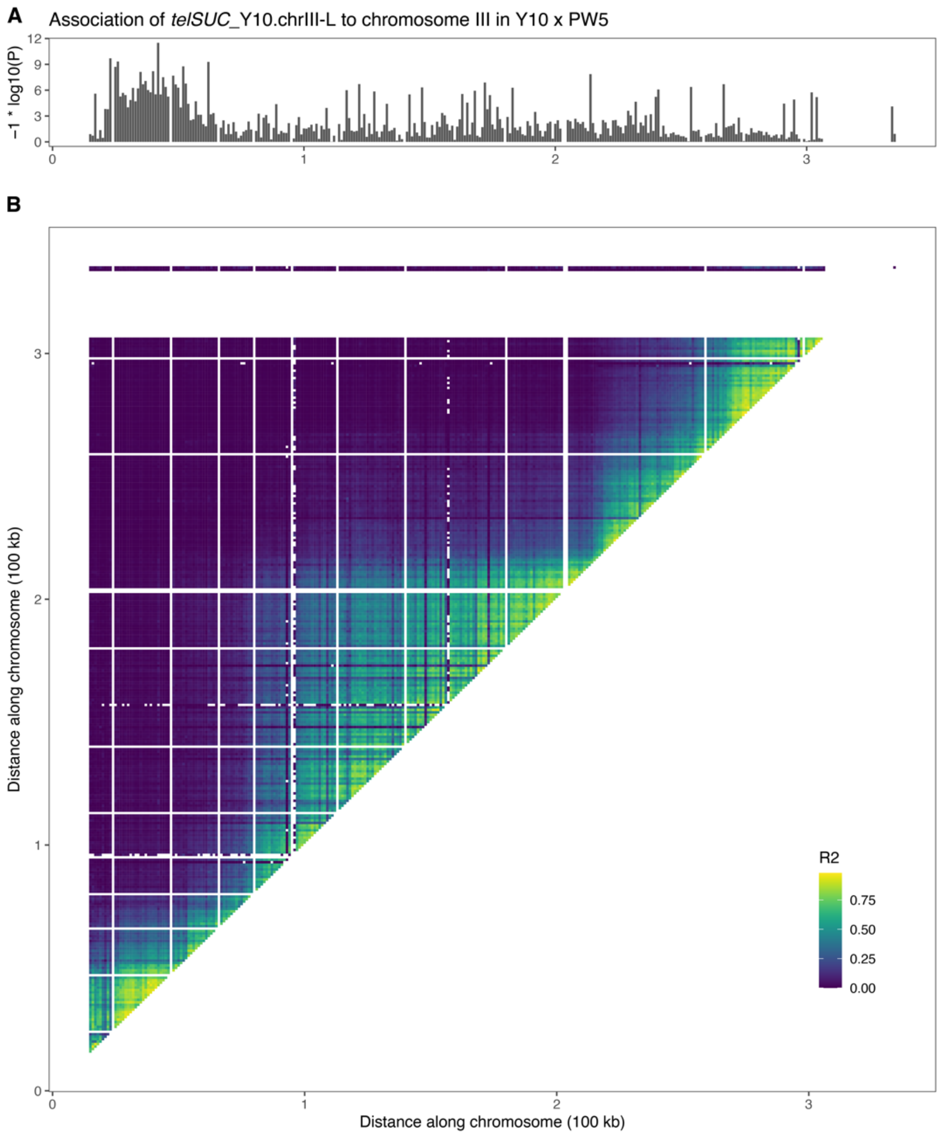


**Figure S12. Validation of assembly of *telSUC*$_{Y10.chrIII-L}$ onto chromosome III-L through linkage mapping in Y10 × PW5.**
**A.** Chromosome III linkage mapping of *telSUC*$_{Y10.chrIII-L}$. Association is shown as in Supplemental Fig. 1A-C.
**B.** Linkage disequilibrium plot of segregating genotypes on chromosome III in Y10 × PW5. Linkage disequilibrium is shown as in Supplemental Fig. 1D.

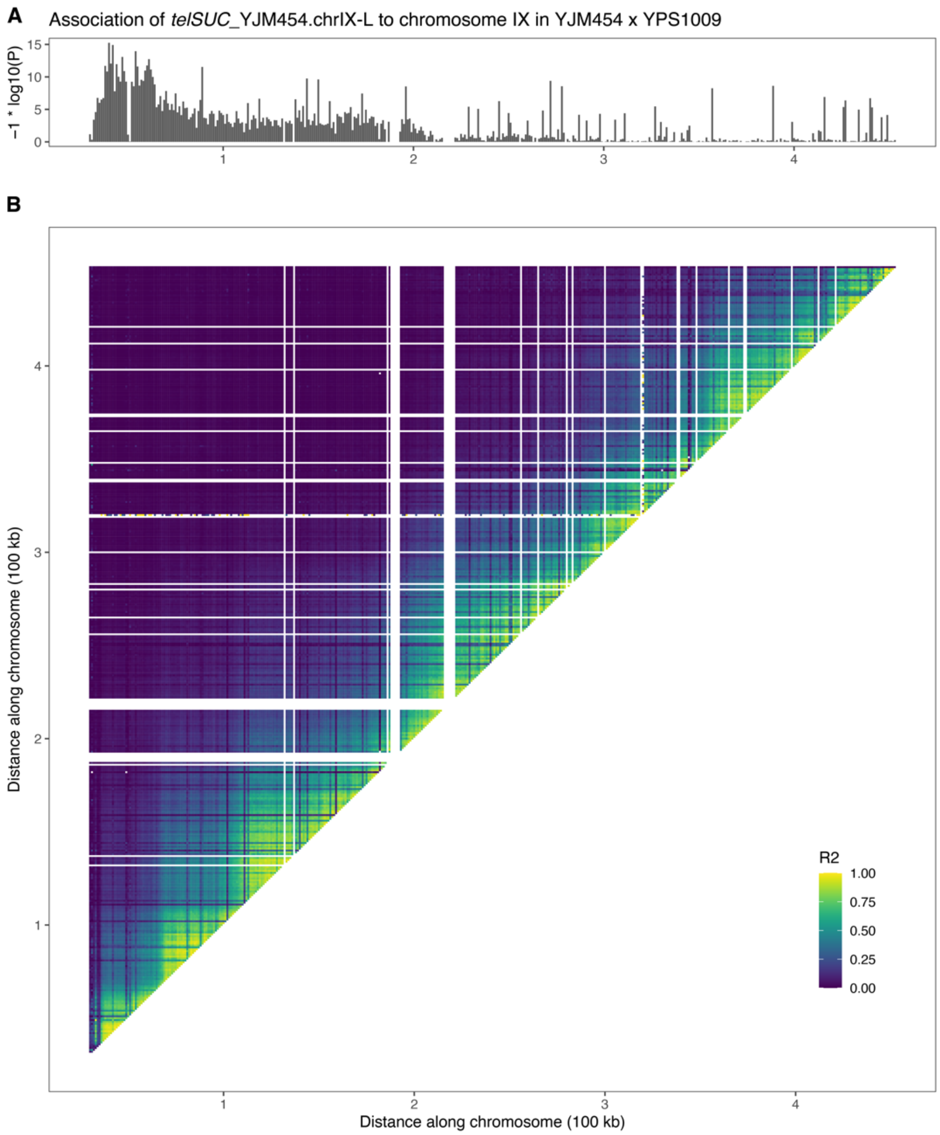


**Figure S13. Validation of assembly of *telSUC*$_{YJM454.chrIX-L}$ onto chromosome IX-L through linkage mapping in YJM454 × YPS1009.**

**A.** Chromosome IX linkage mapping of *telSUC*$_{YJM454.chrIX-L}$. Association is shown as in Supplemental Fig. 1A-C. **B.** Linkage disequilibrium plot of segregating genotypes on chromosome IX in YJM454 × YPS1009. Linkage disequilibrium is shown as in Supplemental Fig. 1D.
